# Eco-Evolutionary Graph Theory: Regular Graphs can act as Amplifiers

**DOI:** 10.64898/2026.10.16.759554

**Authors:** George Berry, Christoph Hauert

**Affiliations:** Department of Mathematics, The University of British Columbia, Vancouver, British Columbia, Canada

**Keywords:** Evolutionary graph theory, Moran process, fixation probability, network population structure, eco-evolutionary dynamics

## Abstract

The Moran model is celebrated for its analytical tractability, yet on network-structured populations it produces contradictory predictions dependent on a seemingly arbitrary choice of birth and death update rule, limiting its biological utility. Here we resolve this problem by decoupling birth and death processes and introducing vacant space. We show that this framework unifies existing models, finding that as the death rate tends to zero the ecological model converges to the death-Birth (dB) Moran process, while the Birth-death (Bd) fixation probability emerges as an upper bound in the limit of large fitness differentials. In this framework random regular and lattice graphs can increase the fixation probability of advantageous mutants and decrease the fixation probability of disadvantageous mutants. They are therefore amplifiers of selection, despite being neutral or suppressing in the Moran model, demonstrating how the introduction of vacant space alters evolutionary outcomes on population structures. This unifying approach is biologically motivated, with results varying continuously via scalar parameters rather than switching between discrete modelling choices. This enables its application in real-world scenarios and offers a better understanding of how population structures and connectivity promote or suppress evolution.

## 1 Introduction

The Moran process is the principal individual-based model for the evolution of populations under constant fitness. Introduced in 1958, it has been used in many applications from the evolution of cancer cell populations (Nowak et al. 2003; Komarova 2006) to the evolution of emotions (Dai et al. 2021) and cloud service providers (Han et al. 2024). It serves as the basis for Evolutionary Graph Theory (Lieberman et al. 2005) and is key to modelling finite populations in Evolutionary Game Theory (Taylor et al. 2004). Since its introduction it has become a fundamental model for understanding stochastic evolutionary dynamics in finite populations. P.A.P. Moran removed the assumption of non-overlapping generations from the Wright-Fisher model (Moran 1958) to create the Moran model, while keeping the assumptions of a fixed population at its carrying capacity and correlated birth and death events. Here we remove the assumption that births and deaths occur in a coupled manner in the traditional Moran model to produce the *decoupled Moran model* and report on its dynamics. We find consistent quantitative and qualitative results that do not require the strict conditions of the Moran process.

In the Moran process a population consists of individuals whose fitnesses depend on their type. Although the process allows for different update mechanisms (Débarre et al. 2014; Jamieson-Lane and Hauert 2015; Yagoobi et al. 2023; Brewster et al. 2025), we focus on the Birth-death (Bd) and death-Birth (dB) processes to illustrate some of its limitations. In the Bd process, an individual is chosen to reproduce proportionally to fitness, then another is chosen to die uniformly at random. In the dB process, the order is reversed: an individual is selected to die, followed by a reproduction event. In both cases, fitness acts on the birth, indicated by the capitalization of the ‘B’. This process continues until only one type remains. In the simplest case, there are two types: residents with fitness *f*_*A*_ and mutants with fitness *f*_*B*_. The process can be represented as a one-dimensional Markov chain, with absorbing states at 0 and *N*, the fixed total population size.

To study evolution in this model, consider an initial setup with one mutant and *N* − 1 residents. Fixation occurs if the mutant type eventually takes over the population and no residents exist. The chance of this happening is a key statistic in the Moran process - the *fixation probability*. Other measures of evolution in the Moran process exist, such as the fixation time (conditional on one type fixating), the absorption time (either type fixating) or the average fitness of the population over time (Sharma and Traulsen 2022). Here we focus on the fixation probability because it is the most widely used measure, allowing for comparison with previous results.

For both Bd and dB formulations, these coupled updates assume a constant population size, mimicking equilibrium at a single unchanging carrying capacity, where any deviation is immediately corrected. These formulations have similar dynamics on a well-mixed (or ‘unstructured’) population (Kaveh et al. 2015) where encounters are random. However, they have vastly different quantitative and qualitative behaviours in networks, limiting the applicability of the results.

Previous work has focused on both these processes, determining the statistical properties for both Bd and dB. This is effective when these properties are similar, as in the unstructured case. By considering the process as a biased random walk with two absorbing states it can be shown that the fixation probabilities in the Bd and dB processes (Kaveh et al. 2015) are

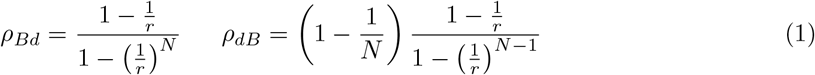

where 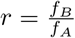 is the ratio between the fitness of the mutant and the resident. The difference between these two fixation probabilities is small and tends to 0 as the population becomes arbitrarily large.

A key difference between the two processes emerges when comparing well-mixed and structured populations which can be represented with networks. These networks can be directed, where edges have a direction and processes can only occur in that direction, or undirected where edges have no direction and processes can occur in either direction. In Bd, as an individual is selected to reproduce first, the competition to reproduce is global and the network may have areas that update more or less quickly (Débarre et al. 2014). In dB, individuals only compete locally and updates happen uniformly at random across the network.

The effects of this distinction are shown when considering the Moran process on undirected regular graphs, where every node has the same in and out degree. In the Bd process, the fixation probability on regular graphs is equivalent to the fixation probability in the well-mixed case, due to the Circulation Theorem (Lieberman et al. 2005). However, this is not the case for the dB process as regular graphs do not have the same fixation probability as in the well-mixed case and the Circulation Theorem does not apply (Kaveh et al. 2015). Figure 1 illustrates this known result, showing that fixation probabilities for regular graphs in the dB process are less than in the well-mixed case.

**Fig. 1:**
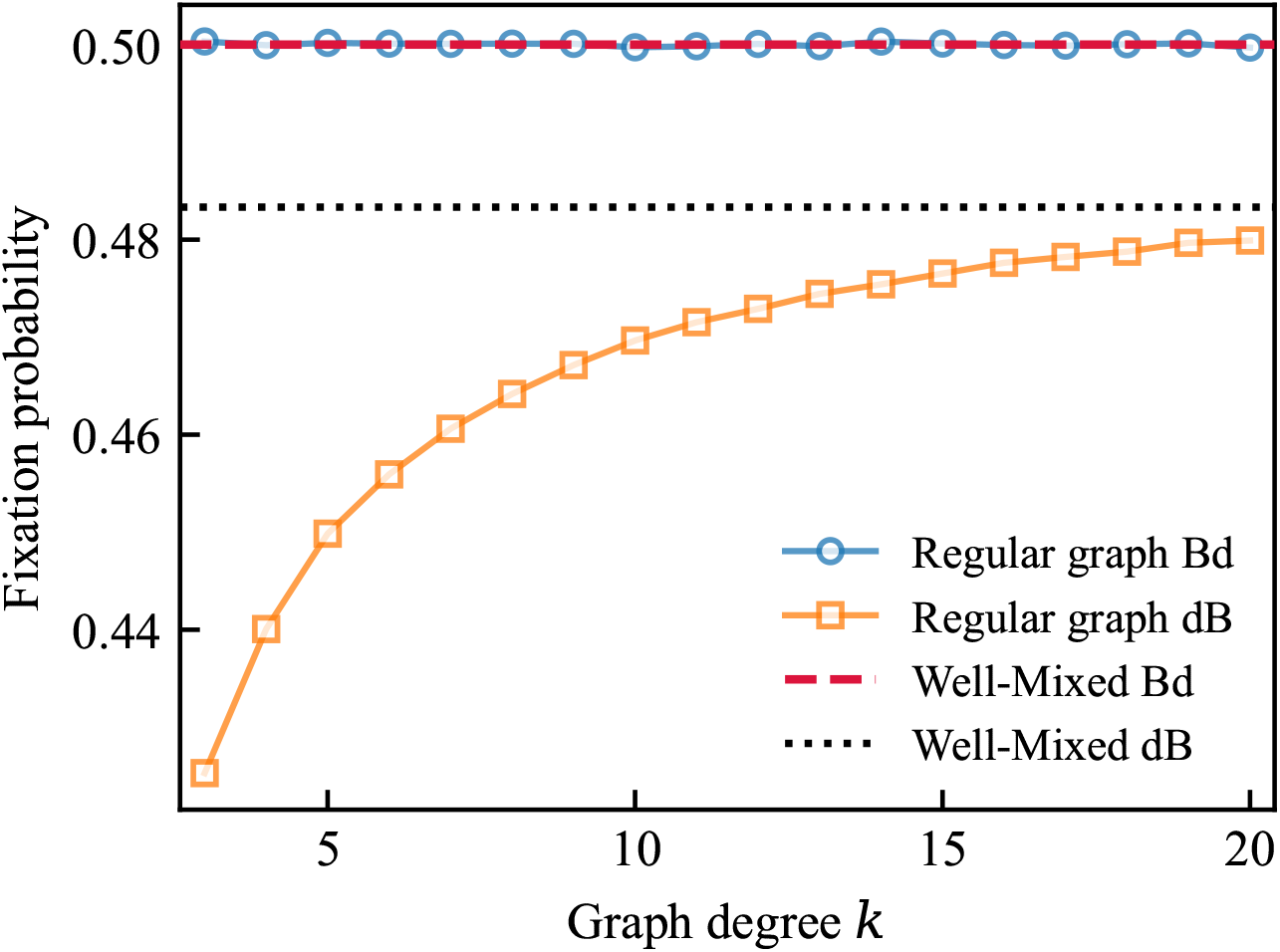
Fixation probabilities in the coupled Moran process as a function of graph degree, comparing Bd and dB updating on well-mixed and regular graphs. The well-mixed lines are based on the known formulas for *ρ*_*Bd*_ and *ρ*_*dB*_ while the regular graph results are determined through simulation. Parameters are *f*_*A*_ = 1, *f*_*B*_ = 2, *N* = 30. In the Bd process, well-mixed and regular graphs fixation probabilities are equal. In the dB process, the fixation probability on regular graphs approaches the well-mixed value from below as the degree tends to *N* − 1. Each point is computed from 10^7^ samples.

The qualitative dynamics become increasingly different when the population has a more complex spatial structure, altering the evolutionary process. We can quantify how they are different by the characterization of amplifiers and suppressors. Amplifiers of selection are population structures in which the fixation probability is greater than in the well-mixed case for beneficial mutants (*r >* 1) and less than the well-mixed case for disadvantageous mutants (*r <* 1). Suppressors of selection are the opposite: they are structures in which the fixation probability is less than in the well-mixed case for beneficial mutants and greater for disadvantageous mutants. For the Bd process the star and directed super-star are amplifiers (Lieberman et al. 2005; Chalub 2016;

Broom and Rychtář 2008). However, for the dB process they are suppressors (Hindersin and Traulsen 2015). Only amplifiers for small fitness differentials have been found for the dB process (Allen et al. 2020), while arbitrarily strong amplifiers have been found for the Bd process on undirected graphs (Pavlogiannis et al. 2018).

These differences in dynamics between the Bd and dB systems make it unclear which update rule most accurately reflects real biological or social processes, limiting confidence in extending the theory to real-world environments. The goal of this work is to find a unifying model that can provide insight into the evolutionary dynamics in ecological populations, determine which results from the dB and Bd systems apply, and give clear indications which population structures will amplify or suppress fixation. Drawing on ecological theory, we meet this goal by decoupling birth and death processes and introducing vacant space, constructing the decoupled Moran process.

## 2 The Decoupled Moran process

To incorporate ecology into the Moran process with minimal modification, consider first a population of only type *A* individuals. Each individual reproduces at rate *f*_*A*_, contingent on vacant space,

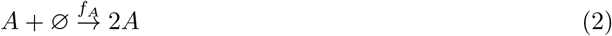

and dies at rate *d*

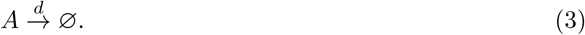

This is the basis of the *decoupled* Moran model, and the original Moran model will be referred to as the *coupled* Moran model for clarity. To determine the dynamics of evolution for this model we must first understand its population dynamics, and the influence that those dynamics have on evolution.

### 2.1 Unstructured Populations

We start by considering the simplest possible scenario: when the population is unstructured, or ‘well-mixed’. Let *i* be the number of individuals and *M* the maximum possible population size in an unstructured population. The population transition rates of the continuous time Markov chain are

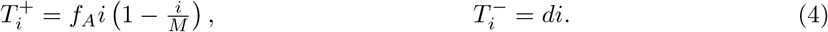

where 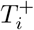 and 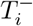 denote the birth and death rates for the whole population, respectively.

The only absorbing state is extinction and the process attains this state almost surely as the population is bounded by *M*. However, we consider the stationary distribution conditioned on non-extinction — the quasi-stationary distribution (Méléard and Villemonais 2012). We define the carrying capacity *K* as the average population size of this quasi-stationary distribution, an emergent quantity determined by the process as in Park et al. (2018). This carrying capacity serves as the ecological equivalent of the fixed population size in the coupled Moran process, allowing for comparisons between the decoupled and coupled models. To understand the quasi-stationary distribution and approximate the carrying capacity, we approximate it using the stochastic differential equation (SDE) (Appendix A) through a system-size expansion (Gillespie 2000):

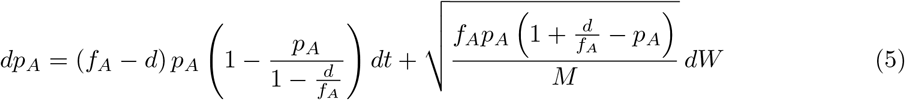

where *p*_*A*_ = *i/M* is the population density, *t* is time and *dW* is the one-dimensional infinitesimal increment of the Wiener process.

In the limit *M* → ∞ the stochastic term vanishes and the SDE reduces to the logistic equation (Verhulst 1838)

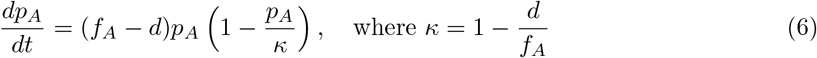

with growth rate *f*_*A*_ − *d* and equilibrium density *κ*. The equilibrium density *κ* can be used to approximate the carrying capacity *K* using *K*_∞_ = *κM*. As *M* increases *K* → *K*_∞_, with Figure 2 showing the distribution skew for small *M* as well as this limit.

**Fig. 2:**
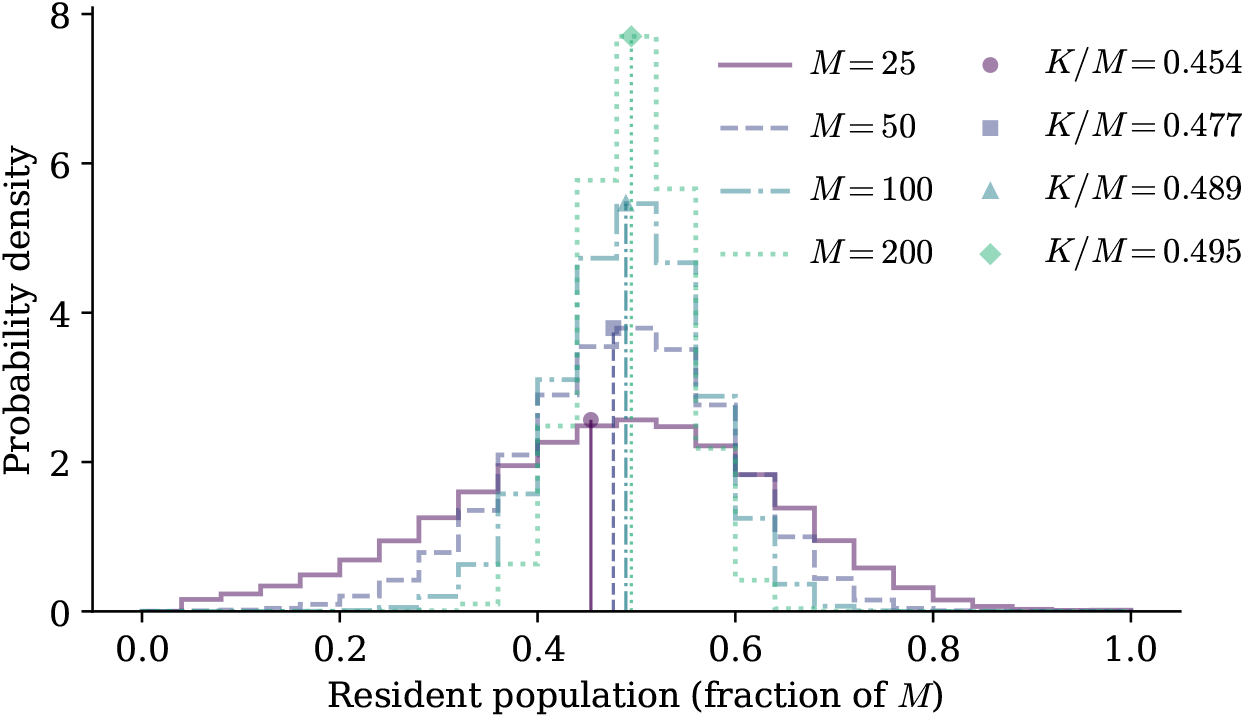
Quasi-stationary distribution histograms of the single-type well-mixed population for different maximum sizes *M*. Parameters are *f*_*A*_ = 1 and *d* = 0.5, for which the deterministic carrying-capacity approximation is *K*_∞_ = 0.5*M*. As *M* → ∞, the empirical carrying capacity approaches *K*_∞_ and the distribution becomes more symmetric with a reduced variance. Vertical lines indicate the empirical carrying-capacity density *K/M* for each *M*, which tend to 0.5.

Mean trajectories of the process demonstrate convergence to *κ* with similar paths from the trajectory mean and logistic equation (Figure 3). As the noise in Equation 5 is multiplicative, stochastic fluctuations generate variance in *p*_*A*_, which shifts the trajectory mean below the deterministic logistic solution. This is a consequence of Jensen’s inequality (Jensen 1906): since the drift in the logistic equation is concave the mean of the trajectory is less than the trajectory of the mean. This difference tends to 0 as *M* → ∞, consistent with the smaller deviation observed for *M* = 100 than *M* = 50.

**Fig. 3:**
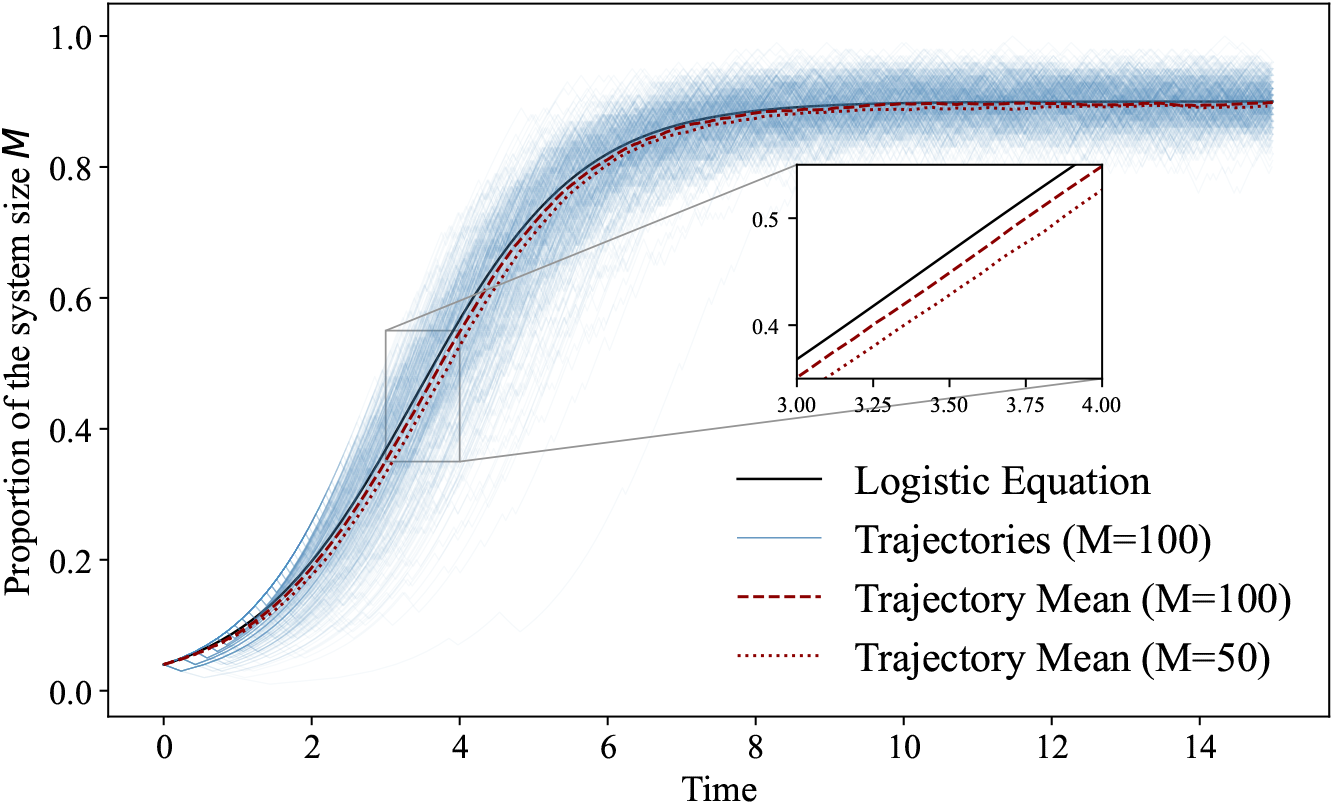
Surviving trajectories of the single-type well-mixed population compared to the logistic equation. Parameters are *f*_*A*_ = 1, *d* = 0.1, and initial condition *p*_*A*_(0) = 0.02, with 500 trajectories used to calculate the mean. The trajectory mean approximately follows the logistic solution, which acts as an upper bound, with larger deviations for *M* = 50 than for *M* = 100. Trajectories are more tightly concentrated around the mean and continue to fluctuate around it even after convergence.

### 2.2 Network Population Structures

In structured populations, each node in the structure is either occupied or vacant, and individuals occupying a node must be adjacent to a vacant space to reproduce. For a node of degree *k* with *v* adjacent vacant nodes, the transition rates are:

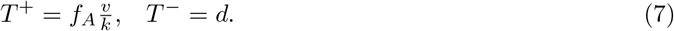

The key quantity is 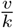, the proportion of adjacent nodes that are vacant. Unlike in the well-mixed case, this varies across nodes, so the population cannot be treated as a single homogeneous unit. To analyze this system we employ pair approximation (Szabó and Fáth 2007; Matsuda et al. 1992; van Baalen 2000), extending mean-field models by explicitly accounting for correlations between neighbouring nodes. Let *p*_*AV*_ denote the proportion of edges connecting an occupied and a vacant space, and define the probability that a neighbour of an individual is vacant as

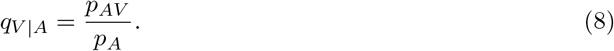

We can approximate *v/k* with *q*_*V* |*A*_ such that an individual reproduces at approximate rate *f*_*A*_*q*_*V* |*A*_. This approximation produces the SDE

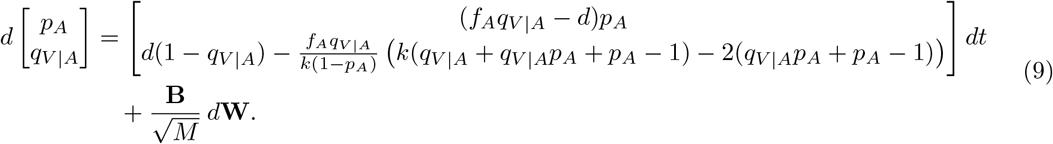

where *k* is the average degree of the network, **B** is the diffusion matrix and **W** is a vector of independent Wiener processes (Appendix A). This approximation ignores higher order connections (e.g. triples like *V* − *V* − *A*) and more complex structures within the graph, but allows for some analytical results. Trajectories of the stochastic process are compared to the deterministic solution in Figure 4 showing close agreement between the two. The behaviour of *p*_*A*_, though similar to the well-mixed population, increases to the equilibrium density more slowly. This is due to the initial local equilibration of the *q*_*V* |*A*_ trajectory before the larger global equilibration.

**Fig. 4:**
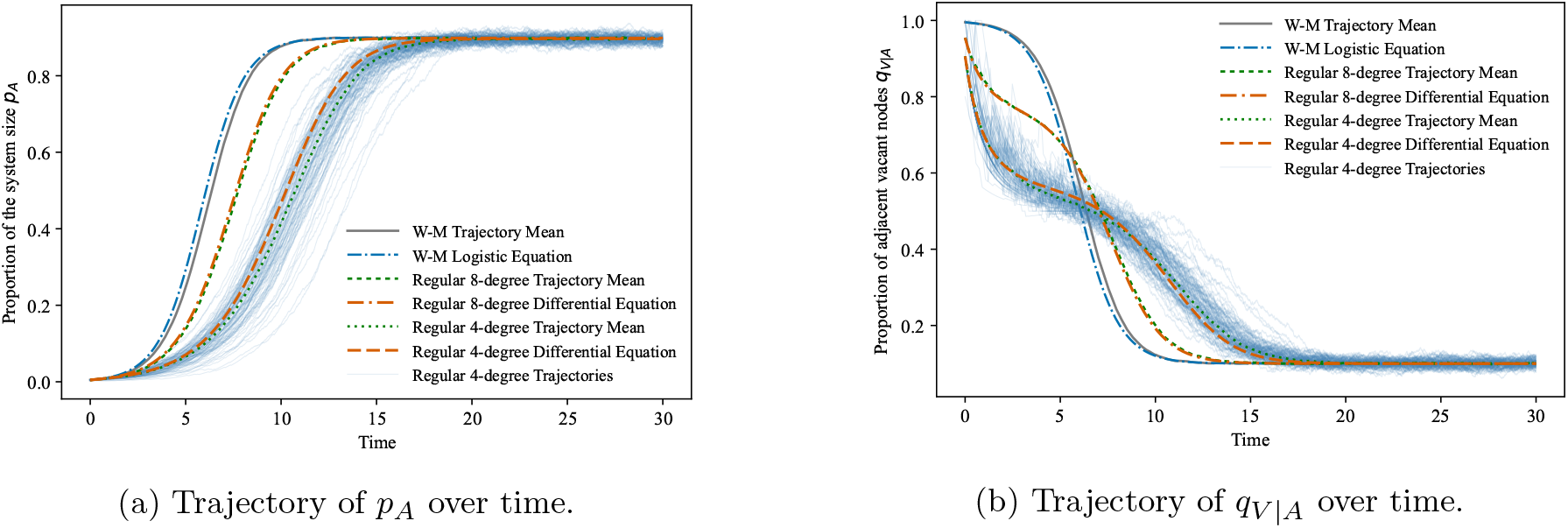
Trajectories of the decoupled model on different regular graphs and the well-mixed population, showing *p*_*A*_ and *q*_*V* |*A*_ over time. Parameters are *p*_*A*_(0) = 0.004, *d* = 0.1, *M* = 1000, *f*_*A*_ = 1, with 500 trajectories used to calculate the mean. Individual trajectories are shown for the 4-regular graph. All mean trajectories track their deterministic solutions closely, though the random regular graphs lag behind the well-mixed population, an effect that grows more pronounced at lower degree. In the random regular graphs, *q*_*V* |*A*_ initially decreases before exhibiting a local equilibration and then converging to the eventual equilibrium. This effect is more pronounced at lower degree and is absent in the well-mixed case. All configurations ultimately converge to approximately the same *p*_*A*_ and *q*_*V* |*A*_ values.

In the limit *M* → ∞, setting 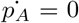 and 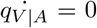 yields the non-trivial equilibrium conditions 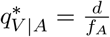 and the approximate equilibrium density for *p*_*A*_:

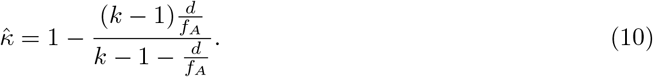

This approximation does not reduce to 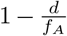, the equilibrium density of the well-mixed population, even when *k* = *M* − 1, which corresponds to the complete graph. Additionally, unlike the well-mixed population, as *M* → ∞ the carrying capacity *K* may not tend to the approximation 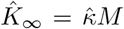. These discrepancies arise from the pair approximation assumption — that we can fully determine the dynamics using just *p*_*A*_ and *q*_*V* |*A*_. These errors vanish in the limit of considering all orders (triplets, quadruplets etc.), which fully encapsulates the state of the graph (Matsuda et al. (1992), Appendix B).

In both unstructured and structured populations, the decoupled model converges to a well-defined quasi-stable distribution, with the carrying capacity *K* playing a similar role to the fixed population size in the coupled Moran process. However, several differences arise from the ecological approach. The population size is no longer fixed but fluctuates stochastically around *K*, introducing demographic noise that is absent in the coupled model. Moreover, extinction is now an absorbing state, though the population’s extinction times are exponential (Ashcroft et al. (2015) and explored in Appendix C). This contrasts with the coupled Moran process, where the population size is fixed and extinction is impossible. Having established these ecological foundations, we now introduce a second type to study evolutionary dynamics and fixation.

## 3 Modelling Evolution in the Decoupled Model

We minimally extend the ecological model by introducing a second type *B* with reproductive fitness *f*_*B*_. The following reaction equations define the decoupled model with both types:

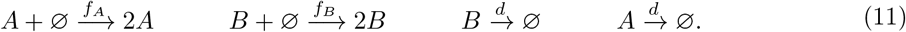

As in the dB and Bd processes, we assume death rates are equal for all types, and evolutionary differences between types are captured solely through their reproductive rates. To motivate the existence of two types, consider *B* as a mutation of the resident *A* (occurring through cosmic rays for example) with mutation rate *µ*:

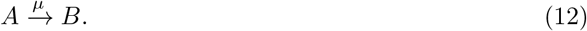

This defines a two-dimensional continuous-time Markov chain, where the state (*i, j*) denotes a population consisting of *i* individuals of type *A* and *j* individuals of type *B*. In the coupled Moran process, the only absorbing state is fixation of one of the types. However in the decoupled process the only absorbing state is extinction of the population so the fixation probability is no longer a suitable term for this setup. Instead we introduce the ‘takeover’ probability. A type is said to ‘take over’ if eventually the population is entirely that type and the other type has died out. We define the takeover probability as the probability that a mutant takes over, starting from a single mutation introduced into a population of residents at its quasi-stationary distribution. To allow for either type to take over before the next mutation we assume that *µM* ≪ 1.

### 3.1 Unstructured Populations

From a state (*i, j*), the possible transitions are births and deaths of either type. The corresponding transition rates in the well-mixed case are

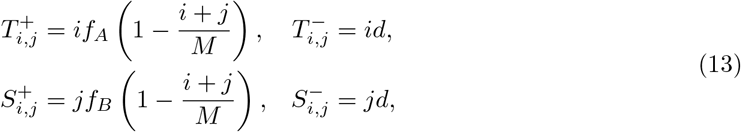

where 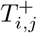 and 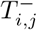 denote, respectively, the birth and death rates in the *A* type, and 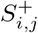 and 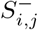 denote the corresponding rates for the *B* type.

As before, let *p*_*A*_ = *i/M* and *p*_*B*_ = *j/M* denote the densities of *A* and *B*. Each type’s drift and diffusion is unchanged from Equation 5 other than the proportion of vacant nodes depends on both types: *p*_*V*_ = 1 − *p*_*A*_ − *p*_*B*_. Therefore the two type SDE is

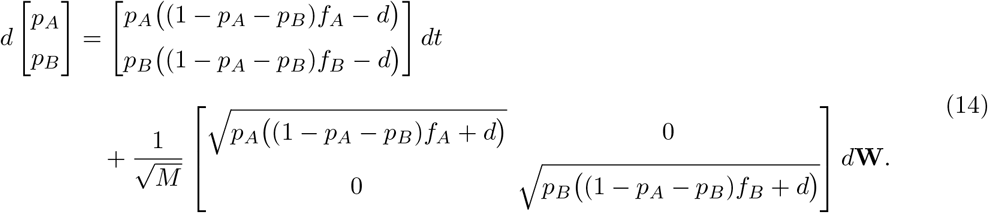

The diagonal form of the diffusion matrix arises from the birth and death reactions only involving one of the two types at a time. The two types never interact directly, competing solely through indirect competition for vacant space. In the limit *M* → ∞, the equilibrium states are

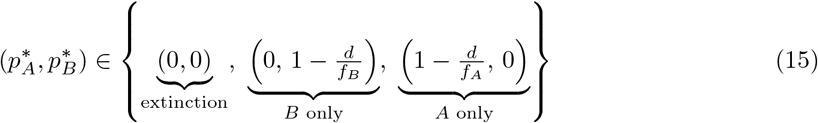

unless *f*_*A*_ = *f*_*B*_ which produces the additional neutral equilibrium state consisting of only random drift, where 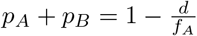.

By defining 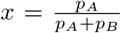 as the proportion of type *A* and *ν* = *p*_*A*_ + *p*_*B*_ as the total density and taking the limit *M* → ∞

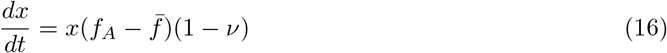

where 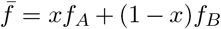 is the average fitness of the population. This corresponds to a non-linear rescaling of time in the standard replicator equation 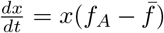 (Taylor and Jonker 1978), with scale factor 1 − *ν >* 0. Since time rescaling preserves orbits and fixed points, their stability and the qualitative phase portrait are identical to those of the unscaled replicator system. The (1 − *ν*) factor captures the intuition that selection requires vacant space in order to act, and determines the speed of the trajectory along each orbit. The dynamics of the total population density in the deterministic limit are given by

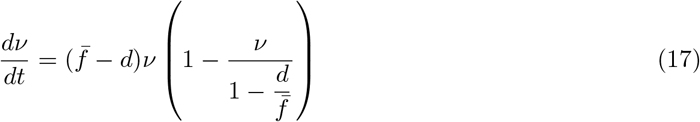

yielding the logistic equation with growth rate 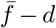 and equilibrium 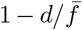. With the evolutionary dynamics established in the well-mixed population, we turn to the methodology required to compare it to other population structures.

### 3.2 Comparing Population Structures

In the coupled Moran process, the fixation probability of a neutral mutant when introduced uniformly at random is 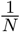 where *N* is the (constant) population size. In the decoupled process, the takeover probability *ρ*_0_ of a neutral mutant depends on the population size when the mutant was introduced. Assuming mutations occur for each individual independently, the total mutation rate scales with population size. Therefore, the population distribution conditioned on a mutation occurring is skewed upwards when compared to the (unconditioned) population distribution, as higher population sizes are overrepresented because they generate mutations at a greater rate. The probability of a mutant being introduced into a population of size *i* is weighted by the probability of the population being at size *i*:

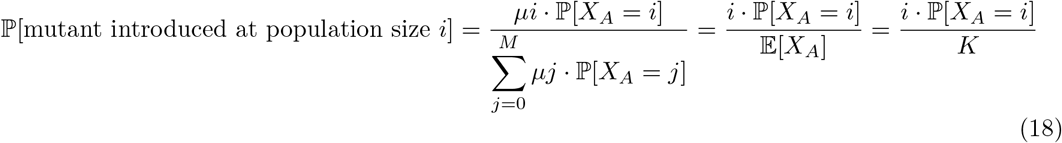

where *X*_*A*_ is the number of residents in the population when the mutant is introduced. Conditioning on the (resident) population size we find that

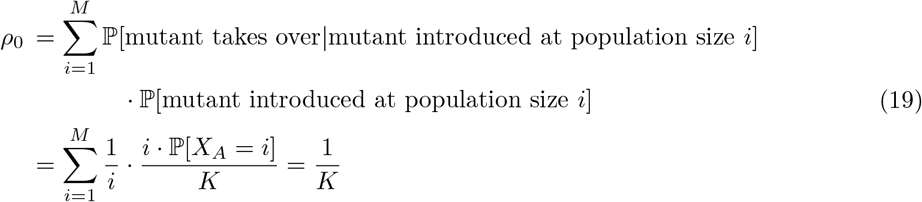

Keeping the neutral takeover probability equal across different configurations is therefore equivalent to keeping *K* equal. This allows for comparisons both with different population structures and also between the coupled and decoupled models by setting *N* = *K*.

To keep *K* constant across different graphs and configurations we vary the graph size *M* and simulate a relaxed resident population using a Gillespie algorithm, conditioning on non-extinction, until the moving average has become approximately constant. Any simulation leading to extinction is discarded. All graphs were configured in this way to allow for comparison with each other and the well-mixed population, setting *K* = 100 where possible - see Figure 5. Without loss of generality the resident fitness is kept constant at 1 (so *r* = *f*_*B*_) and the mutant fitness is varied for death rates *d* = 0.2 and *d* = 0.5. For higher death rates, we approximate the threshold at which the equilibrium is no longer positive by assuming that the approximate equilibrium density 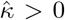, which requires 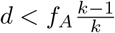. Results for death rates above this threshold are shown in Figure A2.

**Fig. 5:**
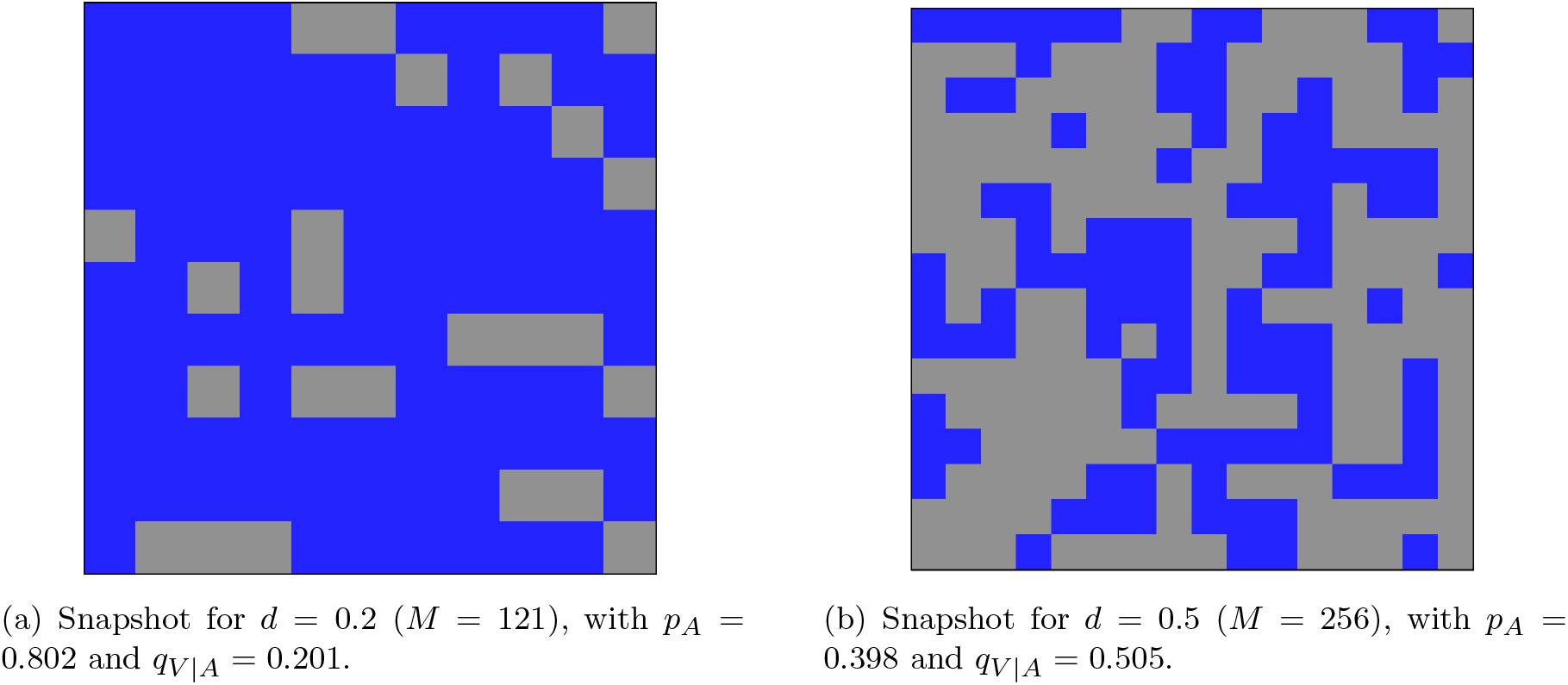
Example populations in a periodic lattice of degree 4 with approximately equal carrying capacities (*K* ≈ 100) but at different death rates and population sizes. The carrying capacities are only approximately equal because of topological requirements of the graph e.g. 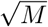 must be an integer for lattice graphs. Blue squares represent inhabited sites and grey squares represent vacant sites. The increased graph size for higher death rates introduces additional vacant sites while keeping the carrying capacity equal.

## 4 Results

### 4.1 Well-Mixed

In the well-mixed population the state is fully described by (*i, j*) and the takeover probability can therefore be obtained from the associated two-dimensional Markov chain, with *D* = (*M* +2)(*M* +1)*/*2 possible states as *i* + *j* ≤ *M*. Let *u*_*i,j*_ denote the takeover probability of the mutants starting with *I* residents and *j* mutants. Then:

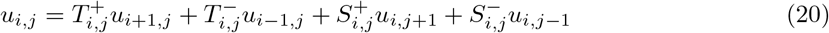

with boundary conditions *u*_0,*j*_ = 1 and *u*_*i*,0_ = 0. Finding takeover probabilities *u*_*i,j*_ therefore becomes equivalent to solving the equation *Q****u*** = *b* where *Q* ∈ ℝ^*D*×*D*^ is the transition matrix, *b* ∈ ℝ^*D*^ is a vector with entries equal to 1 for mutant takeover states and 0 otherwise, and ***u*** ∈ ℝ^*D*^ contains the takeover probabilities across all states. Averaging over the distribution of population sizes at the time of mutant introduction gives the takeover probability

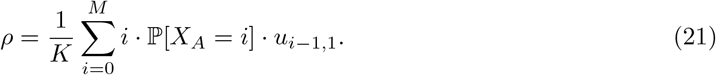

The decoupled process on the well-mixed population produces takeover probabilities similar to the well-mixed population’s fixation probabilities in the coupled Moran process (Figure 6). They converge to the dB fixation probability as *d* → 0 while keeping *r* constant (Figure 7). This is due to the dynamics of the decoupled model tending to the dB process in this limit. Conversely, as the mutant fitness *r* → ∞ the takeover probability approaches a limit close to the fixation probability of the Bd process, which provides an upper bound for the well-mixed case. Note that neither the dynamics nor the takeover results of the decoupled process are tending to Bd process as *r* → ∞. Instead, *ρ*_*Bd*_ → 1 as *r* → ∞, while the decoupled takeover probability tends to a value less than 1. This limit is the probability the mutant has an adjacent vacant site before it dies and is independent of the mutant fitness *r*. This is consistent with analytical results in Appendix E.

**Fig. 6:**
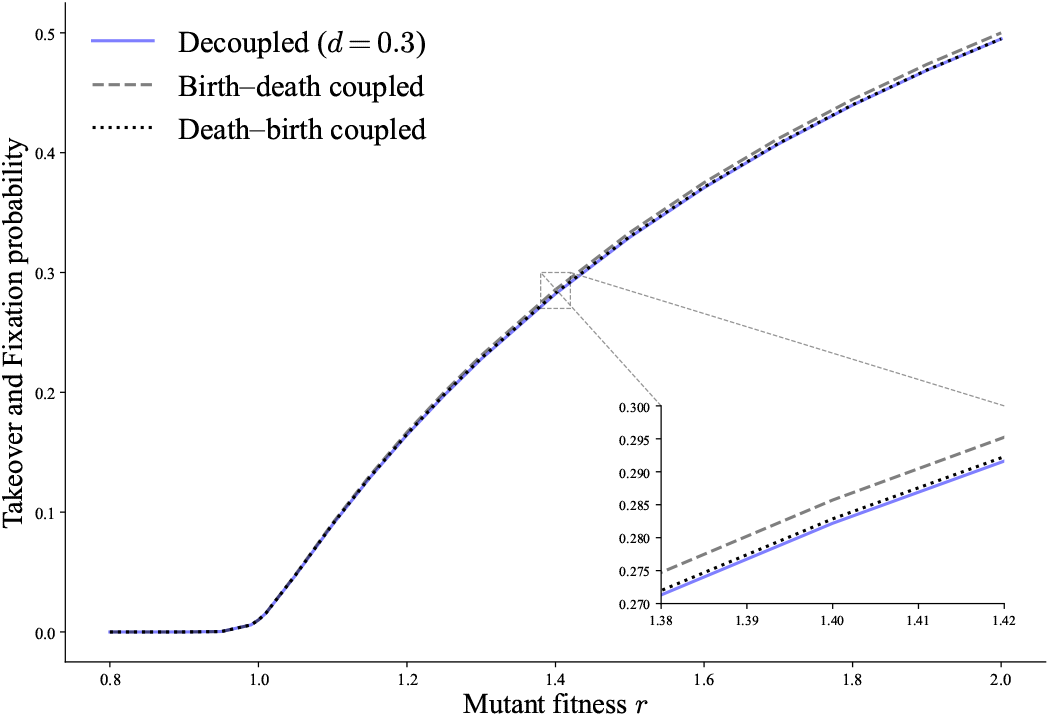
Takeover probabilities as a function of mutant fitness in a well-mixed population, comparing the decoupled process to the Bd and dB coupled Moran processes. Parameters are *N* = 100 in the coupled processes and (*d, M*_*W M*_) = (0.3, 143) in the decoupled process to approximately set *K* = 100. The takeover probabilities of the decoupled process follow the coupled fixation probabilities, and lies above the dB process and below the Bd process for large mutant fitness. Each point is computed from 10^6^ samples.

**Fig. 7:**
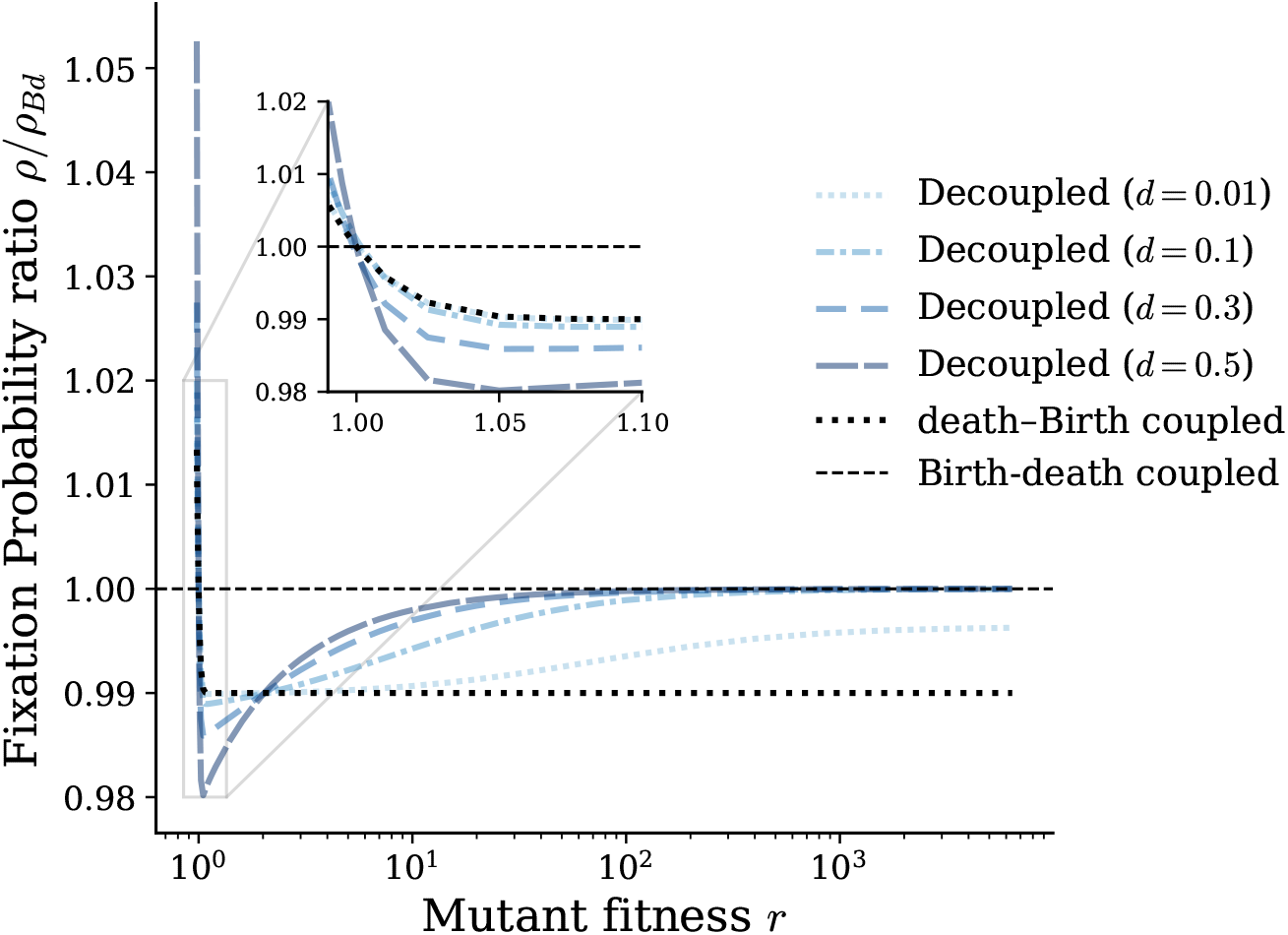
Ratio between takeover (fixation) probabilities *ρ/ρ*_*Bd*_ for the decoupled and dB processes relative to the Bd Moran process. *N* = 100 in the coupled processes and *M* is varied to set *K* = 100 in the decoupled process. (*d, M*_*W M*_) pairs are (0.01, 101), (0.1, 111), (0.3, 143), (0.5, 200). As the death rate decreases the takeover probability tends to the dB fixation probability. For large mutant fitness the takeover probability lies between the dB and Bd fixation probabilities and tends to a constant value for each death rate (Appendix E). Each point is computed from 10^6^ samples.

For the remaining graphs, comparisons are made against the well-mixed population in the decoupled process with equal neutral takeover probability. A graph is said to amplify (respectively suppress) selection at fitness ratio *r* if its takeover probability exceeds (respectively falls below) this baseline. We first consider regular graphs of varying degree, generated either by random edge assignment or by a lattice structure. Random regular graphs isolate the effect of degree distribution, while lattice regular graphs capture spatial constraints and short-range interactions. We then turn to non-homogeneous networks, specifically Barabási–Albert scale-free graphs, whose degree distribution follows a power law, varying the average degree to assess the effect of this non-homogeneity.

### 4.2 Random Regular Graphs

First we consider homogeneous structures, specifically random regular graphs. In contrast to the well-mixed case, takeover probabilities on regular graphs are computed via stochastic simulations in EvoLudo (Hauert 2025) for death rates *d* = 0.2 and *d* = 0.5. Regular graphs exhibit qualitatively similar behaviour to the well-mixed population (Figure 8). However, Figure 9a and Figure 9b, which display ratios relative to the well-mixed baseline, reveal distinct structural effects.

**Fig. 8:**
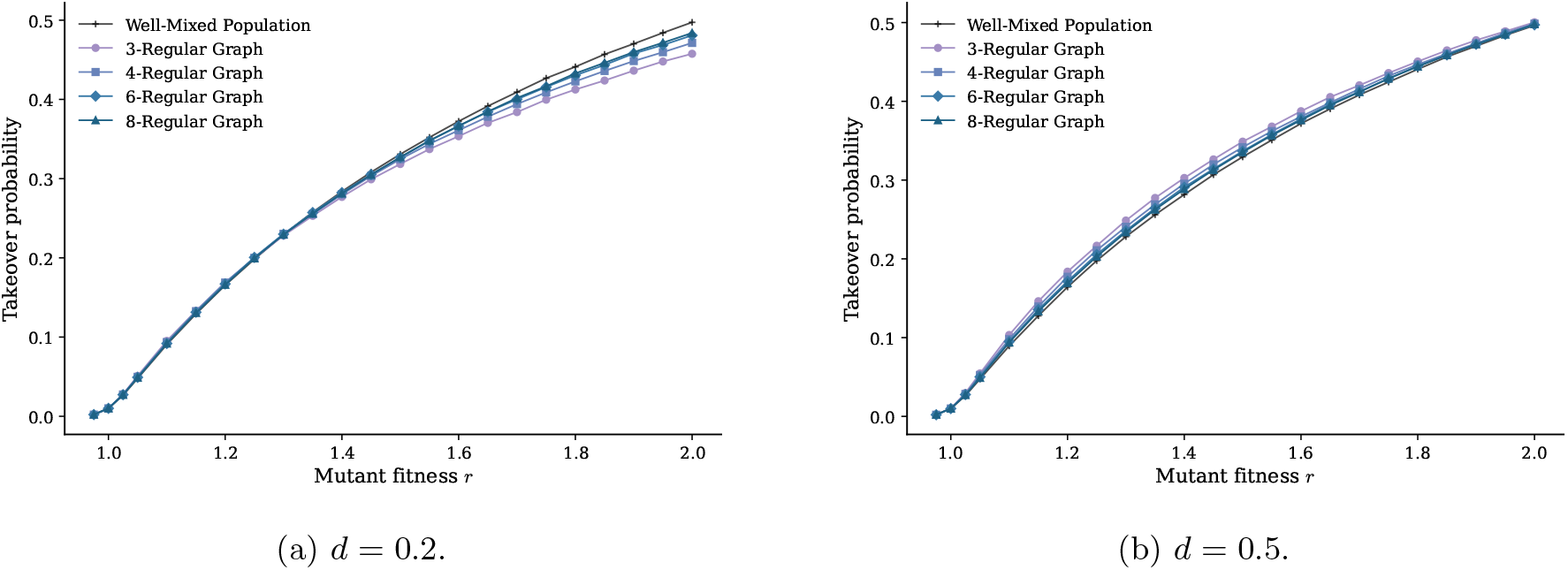
Takeover probabilities as a function of mutant fitness for the decoupled process on random regular graphs, comparing degrees *k* ∈ {3, 4, 6, 8} to the well-mixed baseline. Graph sizes *M* are chosen to approximately set *K* = 100: for *d* = 0.2, *M*_*W M*_ = 125, *M*_*k*=3_ = 129, *M*_*k*=4_ = 128, *M*_*k*=6_ = 127, *M*_*k*=8_ = 126; for *d* = 0.5, *M*_*W M*_ = 200, *M*_*k*=3_ = 341, *M*_*k*=4_ = 259, *M*_*k*=6_ = 230, *M*_*k*=8_ = 219. The takeover probabilities in the regular graphs follow those in the well-mixed population with increasing takeover probability as the mutant fitness increases. Each point is computed from 10^6^ samples.

**Fig. 9:**
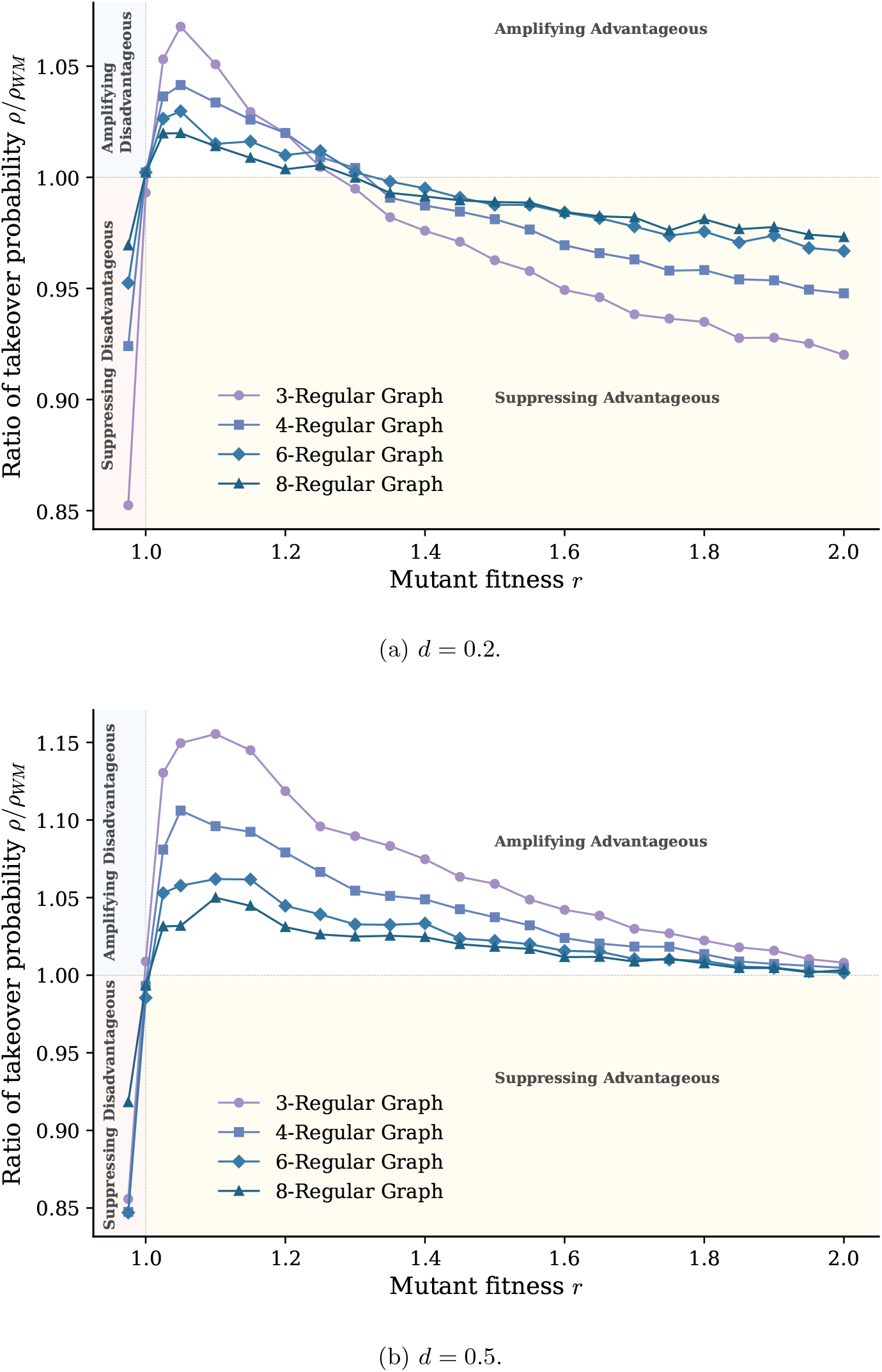
Ratio between takeover probabilities *ρ/ρ*_*W M*_ for the decoupled process on random regular graphs, comparing degrees *k* ∈ {3, 4, 6, 8} to the well-mixed baseline. Graph sizes *M* are chosen to approximately set *K* = 100: for *d* = 0.2, *M*_*W M*_ = 125, *M*_*k*=3_ = 129, *M*_*k*=4_ = 128, *M*_*k*=6_ = 127, *M*_*k*=8_ = 126; for *d* = 0.5, *M*_*W M*_ = 200, *M*_*k*=3_ = 341, *M*_*k*=4_ = 259, *M*_*k*=6_ = 230, *M*_*k*=8_ = 219. Each point is computed from 10^6^ samples. Both death rates show a ratio less than 1 for *r <* 1, passing through *r* = 1, *ρ/ρ*_*W M*_ = 1. This peaks near *r* ≈ 1.05–1.15 with peak height decreasing with degree, followed by a gradual decline as *r* increases. For *d* = 0.2, this decline crosses into suppression by *r* ≈ 1.3, with lower-degree graphs decreasing fastest. For *d* = 0.5, all curves remain amplifying for advantageous mutants throughout the plotted range, converging toward *ρ/ρ*_*W M*_ ≈ 1 by *r* = 2.

For weak selection when *r* = 1+*w* where *w* is small, regular graphs amplify selection: the takeover probability is higher than in the well-mixed population for advantageous mutants and lower for disadvantageous mutants. Additionally, the amplification effect increases as the degree of the graph decreases. It is further increased by increasing the death rate up to the point at which the resident population cannot be sustained in the graph and the carrying capacity is 0 (see Figure A2).

For sufficiently large *r* however, this trend reverses such that regular graphs become suppressors of selection, where the lower the degree of the graph the stronger the suppression. This is consistent with all graphs being suppressors for sufficiently large *r*, as proved in Appendix E and shown in Figure A6, and similarly proved for the coupled dB Moran model in Tkadlec et al. (2020). The transition point between amplification and suppression shifts to larger *r* as the death rate increases.

We attribute the deviation in takeover probability between regular and well-mixed graphs to the availability of vacant nodes, which alters the effective fitness differential between mutant and resident. If the mutant is adjacent to a vacant node it is able to reproduce onto it — it is able to *express* its fitness. If there are no adjacent vacant nodes it cannot reproduce and therefore the effective fitness differential between mutant and residents is 0.

To understand how this concept of effective fitness differential occurs, consider a growing population. Each reproductive event occupies an adjacent vacant node, progressively constraining the birthing individual’s future fecundity. This constraint is more severe at lower degree, since each reproduction occupies a larger fraction of the individual’s neighbouring nodes. We can quantify this effect by approximating the initial expected effective reproductive rates for the mutant using pair approximation. Initially, the reproductive rate of the mutant is 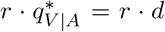, as in Equation 10, which is equal across all graphs under the same death rate. After this initial reproduction, the effective reproductive rate of the resulting mutant pair, calculated using pair approximation, is:

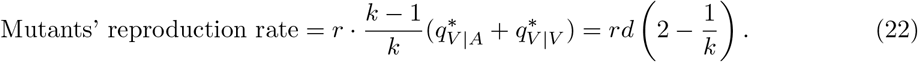

This rate increases with *k*, matching the intuition that lower degrees constrain the growth of a beneficial mutant. As *r* increases, the initial growth and this rate becomes increasingly more important, since the probability of extinction for larger mutant populations decreases. This accounts for the suppression and its varying strength for different regular graphs, as the overall rate of growth is larger in higher degree graphs and maximised in well-mixed populations.

Under weak selection where |*r* − 1| ≪ 1, the decisive quantity is not initial growth but the relative local densities of mutant and resident. Because fitnesses are similar, initially the survivability of the first mutant is similar to that of the surrounding residents. The meaningful differences arise only once the mutant grows to its local equilibrium. After achieving this, for a resident to invade the mutant territory or vice versa there must be vacant nodes for the invader to reproduce into. A high mutant density reduces the number of vacant nodes and reduces the invasive ability of the resident. Likewise a high resident density reduces the invasive ability of the mutant. The relative density governs the relative invasibility of each type, and hence the takeover probability.

In the coupled Moran process, when a resident is replaced by a mutant, the reverse event is symmetric: an invading resident faces the same local environment as an invading mutant. In the decoupled model this symmetry breaks: an invading advantageous mutant in a resident territory encounters more vacant nodes than an invading resident in the mutant territory, since the mutant’s higher carrying capacity leaves fewer vacant sites. The larger this asymmetry, the greater the effect on the takeover probability.

As a metric of that asymmetry, we consider the ratio between the expected density of the mutant and of the resident. To quantify this asymmetry under weak selection, we set *r* = 1 + *w* and approximate the ratio of expected densities with the ratio of equilibrium densities using pair approximation (Appendix B.3):

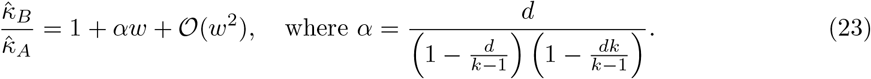

Here 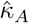 and 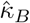 refer to the approximations of the equilibrium density from Equation 10 for type *A* and type *B* respectively and *α* is the equilibrium density ratio slope at *r* = 1. When *w* is sufficiently small *α* governs the size of this ratio. Note that *α* increases as *k* decreases or *d* increases, consistent with the observed amplification behaviour across these parameters. These relationships are supported by simulation results in Figure 10. In the limit *d* → 0 the dynamics converge to a dB process (Appendix E) and consistently *α* → 0, removing the amplification completely and recovering the suppression observed in Figure 1.

**Fig. 10:**
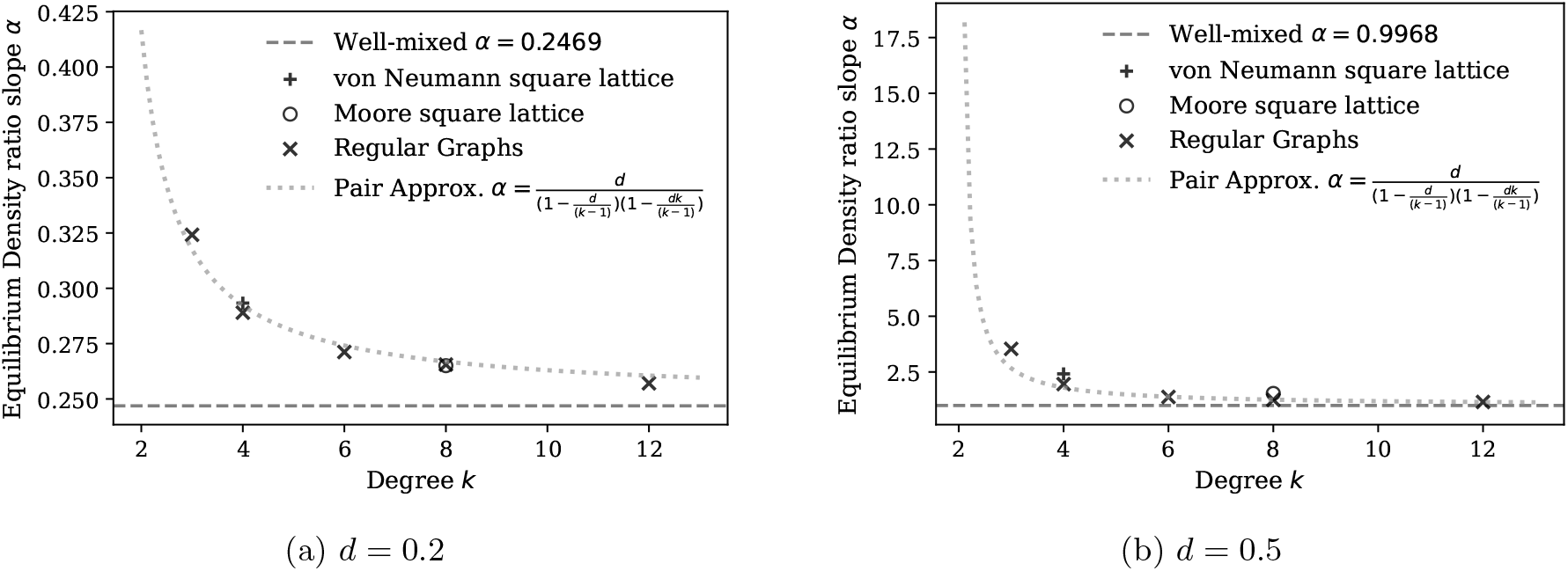
Equilibrium density ratio slope *α* at *r* = 1 as a function of graph degree, comparing random regular graphs to regular lattice graphs. For *d* = 0.2 maximum population sizes are *M*_*W M*_ = 125, *M*_*k*=3_ = 129, *M*_*k*=4_ = 128, *M*_*k*=6_ = 127, *M*_*k*=8_ = 126, *M* = 121 (von Neumann lattice) and *M* = 121 (Moore lattice). For *d* = 0.5 maximum population sizes are *M*_*W M*_ = 200, *M*_*k*=3_ = 341, *M*_*k*=4_ = 259, *M*_*k*=6_ = 230, *M*_*k*=8_ = 219, *M* = 256 (von Neumann lattice) and *M* = 225 (Moore lattice). Lines represent the pair-approximation prediction and well-mixed baseline, and points represent simulation estimates. Random regular graphs follow closely to the pair approximation of *α*, with values decreasing with increased degree and tending to the well-mixed baseline. Lattice graphs exhibit smaller slope ratios than random regular graphs of the same degree.

### 4.3 Regular Lattices

We now examine how spatial constraints and short-range interactions modulate the expected density asymmetry and effective fitness differentials through the Von Neumann lattice (Von Neumann 1966) (4-degree) and Moore lattice (Moore 1962) (8-degree). Both lattices exhibit behaviour qualitatively similar to regular graphs of corresponding degree, but with more pronounced effects (Figure 11). Under weak selection, amplification is stronger than in random regular graphs and for sufficiently large fitness, suppression is likewise stronger.

**Fig. 11:**
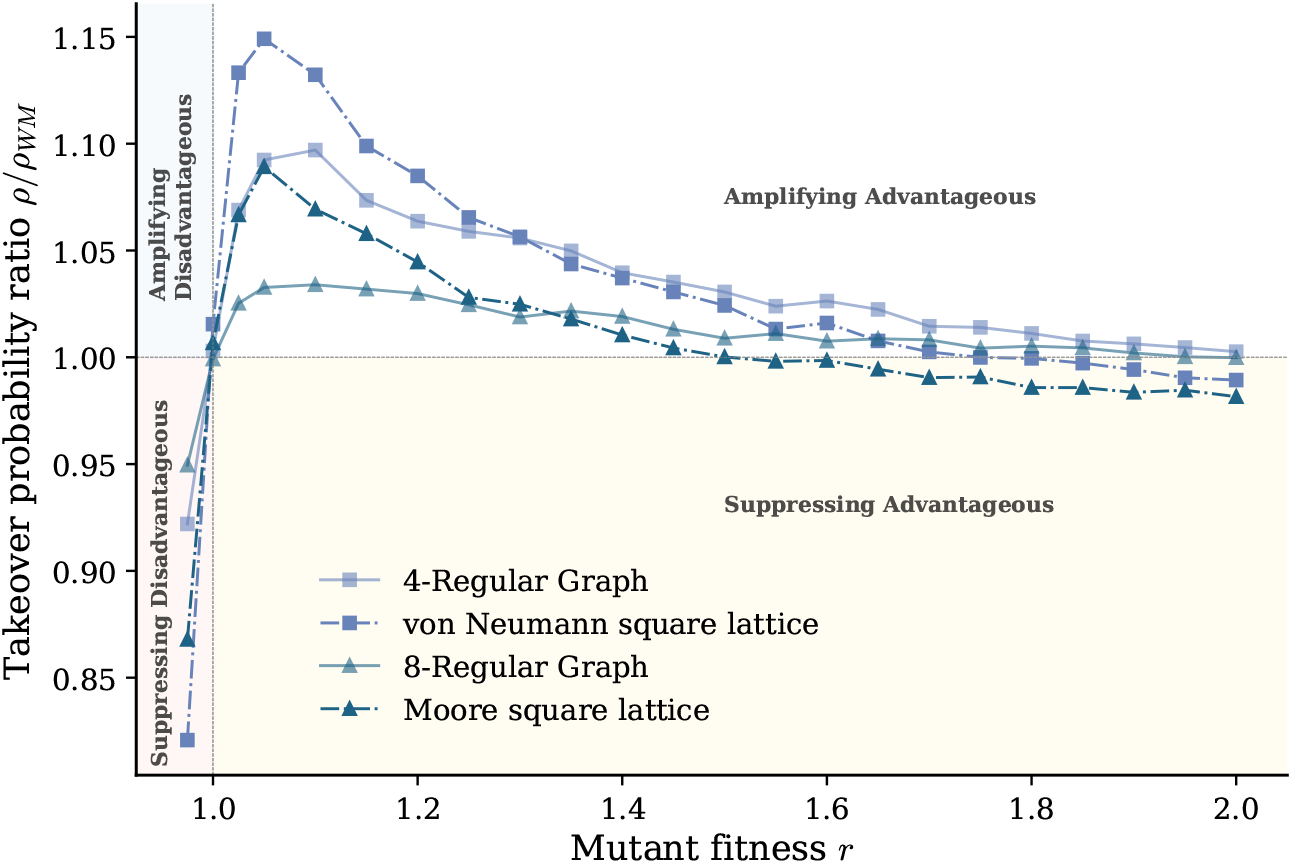
Ratio between takeover probabilities *ρ/ρ*_*W M*_ for the decoupled process on regular lattices, compared to the well-mixed baseline and random regular graphs of the same degree. Parameters are *d* = 0.5 with *M* = 256 (von Neumann lattice) and *M* = 225 (Moore lattice), and *M*_*k*=4_ = 259 and *M*_*k*=8_ = 219 for the corresponding random regular graphs. Each point is computed from 10^6^ samples. All four structures show a ratio below 1 for *r <* 1, a sharp jump to amplification just above *r* = 1, and a peak near *r* ≈ 1.05–1.10 before declining toward *ρ/ρ*_*W M*_ ≈ 1. Each lattice initially amplifies more strongly than its corresponding random regular graph, but declines faster and eventually falls below it, after which the two decline in near-parallel fashion. The lower-degree structures (4-regular, von Neumann) amplify more strongly throughout than the higher-degree structures (8-regular, Moore), though all four converge to approximately equal ratios by *r* ≈ 2.

The expected density asymmetry that drives amplification on regular graphs extends to lattice graphs, though with important differences. Lattice graphs exhibit a higher expected density ratio slope than their regular graph counterparts (Figure 10), which accounts for the increased amplification of weak selection observed in these structures. This distinction is more pronounced at *d* = 0.5 than at *d* = 0.2 (Figure 10a, Figure 10b), consistent with the similar amplification between lattice and regular graphs at *d* = 0.2 (Figure A5) and the divergence in amplification at *d* = 0.5. The stronger suppression effect at large *r* in lattice graphs can be attributed to the cycles inherent to their structure. As growth is more localised, spatial clustering confines the mutant population, reducing its effective reproductive rate following initial expansion and increasing the likelihood of local extinction. This localisation increases the suppression further than the effect of the graph degree, resulting in a lower takeover probability than their corresponding regular graphs.

Both random regular and regular lattice graphs exhibit amplification under weak selection, a novel phenomenon arising from the distribution of vacant sites and the resulting dependence of expected density on fitness. The introduction of spatial structure in lattices boosts this amplification and shifts the onset of suppression (which all graph structures eventually display as *r* → ∞) to lower values of *r*.

### 4.4 Scale-free graphs

Unlike lattices, which impose uniform local structure and short-range interactions, scale-free graphs introduce substantial degree heterogeneity, providing a contrasting regime to test the generality of the carrying-capacity asymmetry in Equation 23.

Barabási-Albert scale-free graphs (Barabási and Albert 1999) are constructed via preferential attachment, where each new edge connects to an existing node with probability proportional to that node’s current degree. This produces a few high-degree hubs and many low-degree nodes, resulting in the power law distribution of degree. Because the graphs used here have fewer than 10^3^ nodes, this scale-free structure is only approximate, but the resulting degree distribution will still be highly heterogeneous. The average degree ⟨*k*⟩ is varied across these graphs to test whether the degree-dependent amplification and suppression effects seen in subsection 4.2 carry over to scale-free graphs.

Figure 12a shows a slightly different behaviour from the regular graphs for low death rates. For weak selection at low death rates there is little to no amplification when compared to the well-mixed population. Instead, the takeover probability is suppressed throughout and as in lattice and regular graphs the suppression increases as the fitness *r* increases. For larger death rates such as in Figure 12b the behaviour is similar to regular graphs with amplification for weak selection, but the graphs suppress at a lower fitness value than in regular graphs.

**Fig. 12:**
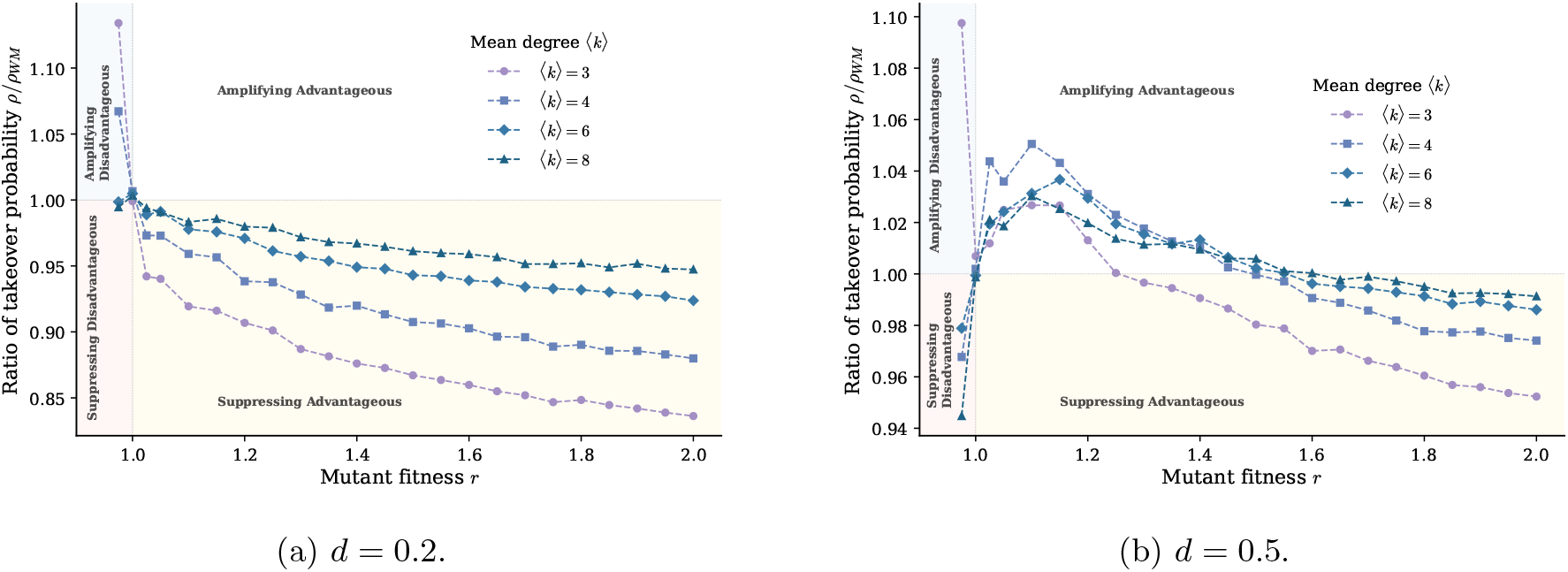
Ratio between takeover probabilities *ρ/ρ*_*W M*_ for the decoupled process on Barabási–Albert scale-free graphs with different average degrees, compared to the well-mixed baseline. Graph sizes *M* are chosen to approximately set *K* = 100: for *d* = 0.2, *M*_*W M*_ = 125, *M*_⟨3⟩_ = 187, *M*_⟨4⟩_ = 160, *M*_⟨6⟩_ = 148, *M*_⟨8⟩_ = 143; for *d* = 0.5, *M*_*W M*_ = 200, *M*_⟨3⟩_ = 811, *M*_⟨4⟩_ = 489, *M*_⟨6⟩_ = 355, *M*_⟨8⟩_ = 313. Each point is computed from 10^6^ samples. For *d* = 0.2, when *r <* 1 all mean degrees show a ratio above 1 (amplifying disadvantageous), most strongly for ⟨*k*⟩ = 3 (*ρ/ρ*_*W M*_ ≈ 1.13), before dropping sharply through *ρ/ρ*_*W M*_ = 1 at *r* = 1. For *r >* 1, the ratio falls immediately below 1 and declines as *r* increases. Lower mean degree produces stronger suppression throughout. For *d* = 0.5, when *r <* 1 the ratio is above 1 for ⟨*k*⟩ = 3 but below 1 for all other degrees. For *r >* 1, all curves briefly amplify, peaking near *r* ≈ 1.05–1.15, before declining as *r* increases. This decline crosses into suppression by *r* ≈ 1.5–1.6, after which lower mean degree again yields stronger suppression.

In the coupled Moran process scale-free graphs are amplifiers of selection under Bd (Lieberman et al. 2005) for all *r*, but this is not the case for the decoupled model. In scale-free graphs, there is no amplification when |*r* − 1| ≪ 1 at low death rates, in contrast to the behaviour observed in homogeneous graphs. This can be attributed to the distribution of the quasi-stationary resident population across nodes of varying degree. Unlike regular and lattice graphs which exhibit homogeneous degree distributions, the resident population in scale-free graphs is concentrated at higher degree nodes, leaving vacant nodes predominantly with lower degree. This is consistent with intuition: a vacant node of higher degree has more occupied neighbours that could reproduce into it. Consequently, the vacant nodes adjacent to an initial mutant are more likely to be of lower degree. Furthermore, the local environment of the mutant differs substantially from the global structure of the graph, such that the expected density ratio is no longer a reliable measure. Since the mutant is restricted to reproducing onto lower-degree nodes, its capacity to spread is diminished relative to a well-mixed population. This is analogous to taking the limit *d* → 0 in which the dynamics approach those of the dB process. Here the topology of the graph accelerates this convergence by concentrating reproduction onto lower-degree nodes. As the death rate increases, the disparity in node occupancy becomes more pronounced, as depicted in Figure 13. Although the resident population occupies very high degree nodes in Figure 13b, the vacant nodes retain sufficiently high degree to permit mutant spread, recovering the amplified dynamics observed in other graph structures, albeit with a weaker amplification effect. This disparity in node occupancy is more pronounced for graphs with low average degree (Appendix B.4). Overall, the degree heterogeneity of scale-free graphs renders residents more resistant to invasion, as the higher degree nodes remain occupied by residents, limiting the mutant’s access to the majority of the graph.

**Fig. 13:**
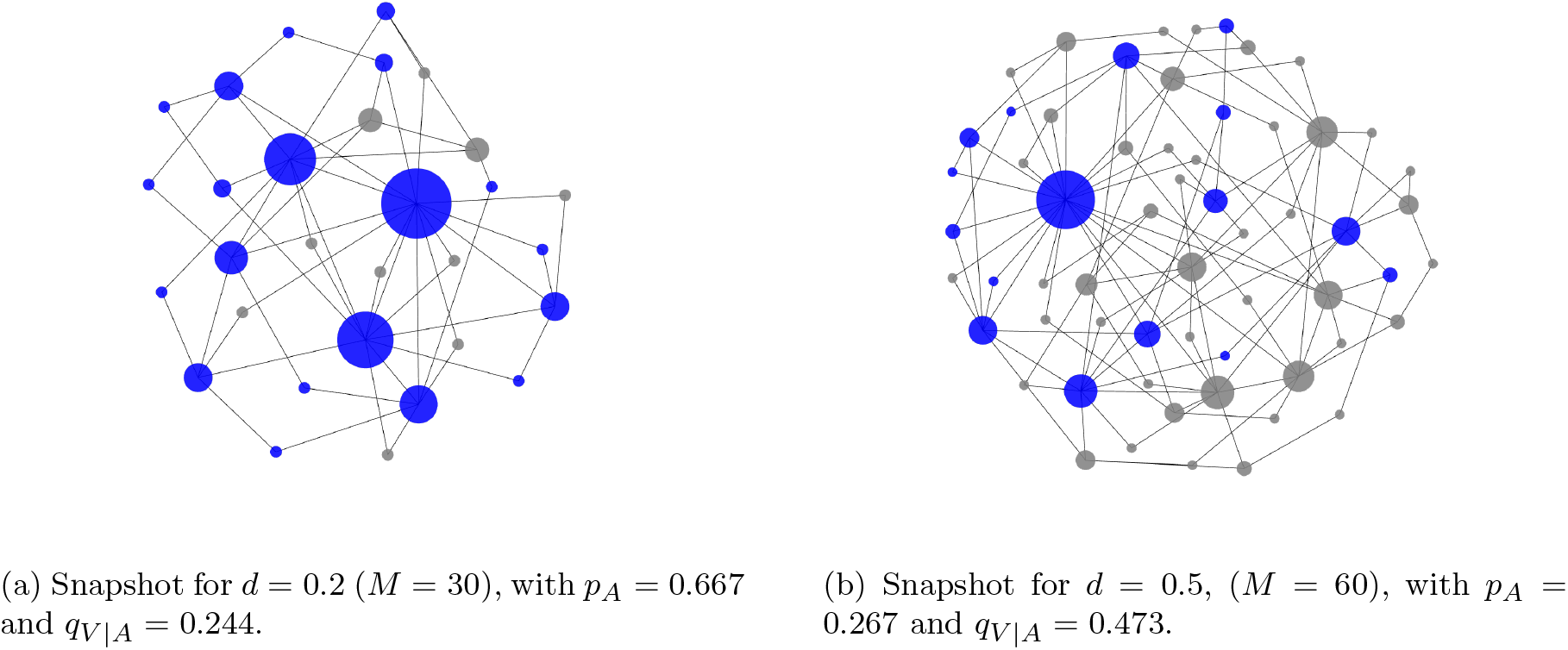
Example populations on a Barabási–Albert graph with ⟨*k*⟩ = 4 and approximately matched carrying capacities (*K* ≈ 20) at different death rates, achieved by varying the graph size *M*. Blue nodes are occupied and grey nodes are vacant, with node size proportional to degree. Higher-degree nodes are occupied more frequently, and lower death rates produce fewer *V* –*V* pairs.

The structure-dependent effects observed in regular, lattice and scale-free graphs illustrate that evolutionary outcomes in structured populations depend significantly on the death rate and graph topology through the distribution of vacant sites, rather than merely on the update-rule conventions of the coupled Moran process. Together these effects produce suppression as *r* → ∞ and the novel phenomenon of amplification under weak selection in regular graphs. The degree heterogeneity of scale-free graphs concentrates residents at high-degree nodes, rendering invasion more difficult and weakening any amplification relative to more homogeneous structures.

## 5 Conclusion

This work explores the decoupled Moran process as an alternative to the coupled Moran process to create a unifying individual-based and ecologically motivated evolutionary framework for network-structured populations. By treating births and deaths as independent processes, the decoupled model resolves the ambiguity in the coupled model of choosing an update rule.

This decoupled approach comes with its own challenges. One of these is that comparing different population structures is more difficult. As the carrying capacity can only be approximated, the graph size must be varied to ensure that the neutral takeover probability is approximately constant (subsection 3.2). The approximation is made worse by the discrete nature of the graphs and the constraints of some graph geometries e.g. square lattices (Figure 5). In these latter cases the target carrying capacity was changed to allow for the comparison. Another challenge is the reduction of analytic tractability, leading to no closed-form fixation results for network-structured populations and only a numerical solution for the well-mixed process (subsection 4.1), as the difference equation is now 2-dimensional. This loss of tractability stems from the introduction of vacant space into the model which also has significant consequences for the evolutionary outcomes.

Takeover results for the well-mixed population are similar to the original coupled Moran process, with the takeover probabilities tending to the fixation probabilities of the dB process as *d* → 0 (Appendix E.1). This limit holds on all network population structures, as the carrying capacity increases to the maximum population size, resulting in the dynamics mimicking the dB process. Additionally, the takeover probabilities are bounded above by the fixation probabilities of the Bd process for large *r* (Appendix E.2). This bound also holds on all network population structures. As *r* → ∞, takeover probabilities tend towards the probability that the mutant has an adjacent vacant site before it dies. As *K* → ∞, this probability tends to 1 for the well-mixed population but remains below 1 for graphs of finite degree. Consequently, takeover probabilities for large *r* are lower on networks than in the well-mixed population, particularly for lower-degree graphs (Figure 9, Figure 11, Figure 12).

A novel result occurs under weak selection due to the introduction of vacant space. Homogeneous graphs, including random regular and regular lattice graphs, can act as amplifiers of selection in this regime (subsection 4.2, subsection 4.3). Previous results for regular graphs showed that they are neutral in the Bd process (Lieberman et al. 2005) and suppressors in the dB process (Hindersin and Traulsen 2015). Heterogeneous graphs can also act as amplifiers (subsection 4.4), consistent with the amplification on scale-free graphs previously reported for the Bd process (Lieberman et al. 2005). This behaviour in both graph classes arises from asymmetric effects on the equilibria of mutants and residents, altering their relative abilities to invade one another. Though amplifiers of selection have been found in the dB process under weak selection (Allen et al. 2020), these amplifiers do not include regular graphs, and do not apply to all graphs as the amplification reported here does. This selective amplification in the decoupled model is of particular note because it applies to those advantageous mutants that need it the most: very advantageous mutants with high fitness are likely to take over regardless, while mutants with *r* close to 1 are generally more subject to random drift.

Overall, the framework provides a natural, ecologically motivated extension of evolutionary graph theory, introducing new dynamics and evolutionary outcomes while remaining free from the ambiguous artefacts of coupled updating. This opens directions for studying new amplifiers and suppressors, alternative mutation initialisations such as through reproduction or migration, and extended metrics of evolution such as takeover times.

## Acknowledgements

We acknowledge financial support from the Natural Sciences and Engineering Research Council of Canada (NSERC) RGPIN-2021-02608.

## A Stochastic Differential Equation Derivations

### A.1 Unstructured Population

The logistic equation for the single-type population in the decoupled model is derived as follows. Consider a single timestepΔ*t* and just the resident population. Let *P* (*t, i*) = P(there are *i* residents at time *t*), which describes the state of the system. We consider the state a single timestep after *t* (at *t* +Δ*t*) to produce the forward Kolmogorov equation for *P* (*t* +Δ*t, i*):

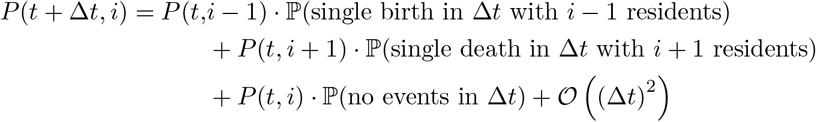

With 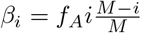 and *δ*_*i*_ = *di* as the total rates of births and deaths respectively,

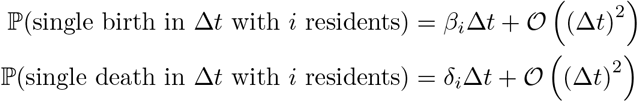

We can then simplify the forward equation to the following.

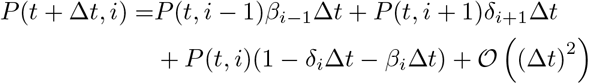

Rearranging this equation gives

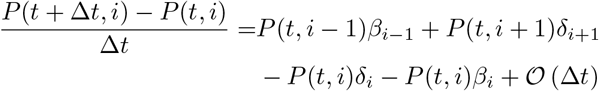

We rescale the resident population by the total size *M*, replacing the variable *i* with *p*_*A*_ where 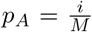 as in the Kramers-Moyal expansion (Gardiner 2009). With this rescaling the change from a single birth or death becomesΔ*p*_*A*_ = 1*/M* rather than 1. To simplify the equations we introduce new variables *µ*_*b*_(*p*_*A*_) = *f*_*A*_*p*_*A*_(1 − *p*_*A*_) and *µ*_*d*_(*p*_*A*_) = *p*_*A*_*d* so that

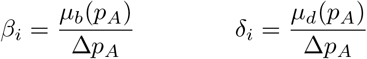

To convert the probability mass function to a probability density function let Π(*t, p*_*A*_) = *M* · *P* (*t, i*). Including these new variables produces:

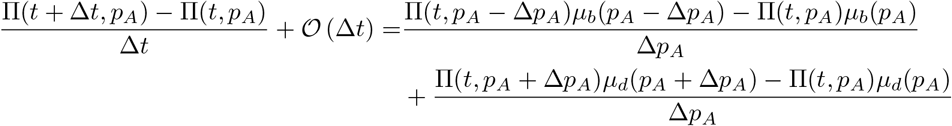

We approximate the left hand side with 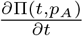 as in Traulsen et al. (2005) & Traulsen et al. (2006). Taking Taylor series approximations around *t* and *p*_*A*_, we can write this as

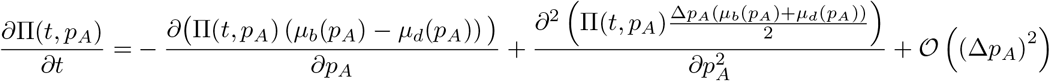

which is the Fokker-Planck equation with error term *O* (Δ*p*_*A*_)^2^ = *O M* ^−2^. Taking the limit *M* → ∞, this yields the stochastic differential equation according to Itô’s Lemma (Itô 1957):

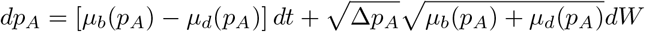

where *dW* is the one-dimensional infinitesimal increment of the Wiener process. We can simplify further by substituting in for *µ*_*b*_, *µ*_*d*_ andΔ*p*_*A*_:

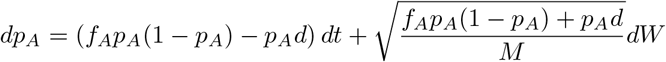

Finally, letting *M* → ∞ the Brownian motion term becomes negligible and the logistic equation is derived

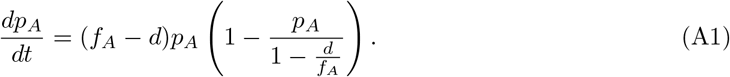

Note that the diffusion coefficient in the SDE is proportional to 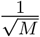, allowing us to quantify the stochastic nature of the system.

### A.2 Network Population Structures

An approximate SDE of *k*-degree random regular graphs can be found using pair approximation and following the same steps. Let *P* (*t, i, j*) = ℙ (there are *i* residents and *j AV* edges at time *t*), which describes the state of the system. The forward Kolmogorov equation for *P* (*t* +Δ*t, i, j*) is

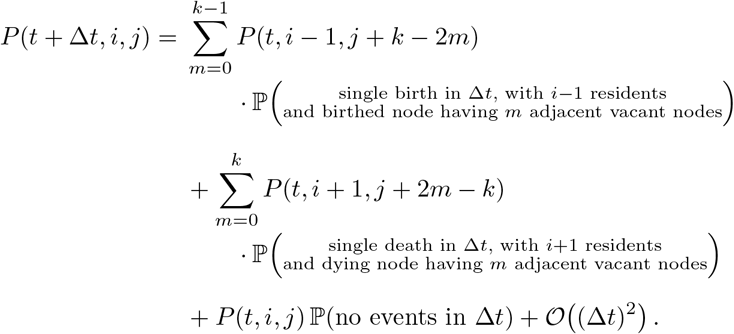

With 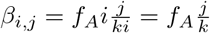 *δ*_*i*_ = *di* as the total rates of births and deaths respectively, we estimate the probabilities using pair approximation

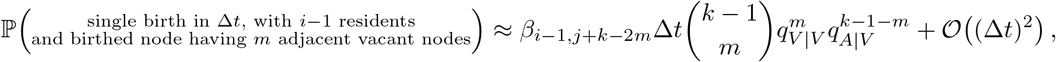

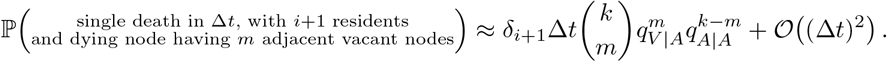

where

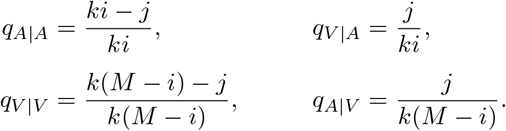

Substituting these values into the master equation gives

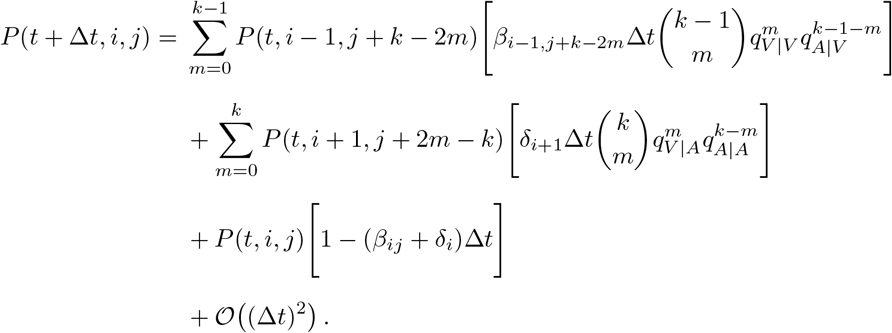

As before, we apply the Kramers-Moyal and Taylor expansions with *p*_*A*_ = *i/M* and *q*_*V* |*A*_ = *p*_*AV*_ */p*_*A*_ and convert the probability mass function to a probability density function,

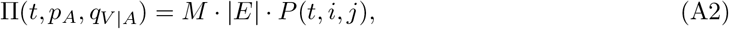

where |*E*| is the number of edges in the graph. Collecting state variables in the vector **x** = (*p*_*A*_, *q*_*V* |*A*_)^⊤^ and writing 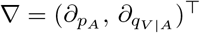 withΔ*p*_*A*_ = 1*/M*, this gives the Fokker-Planck equation

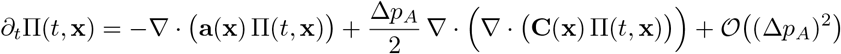

Here **a** is the drift vector and **C** is the symmetric noise matrix,

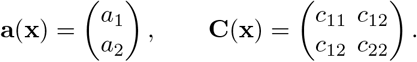

The elements of the drift vector are

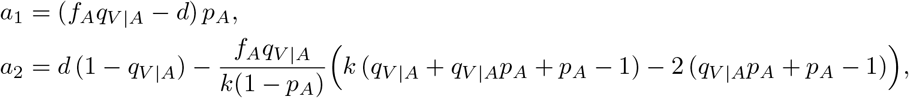

and the elements of the noise matrix are

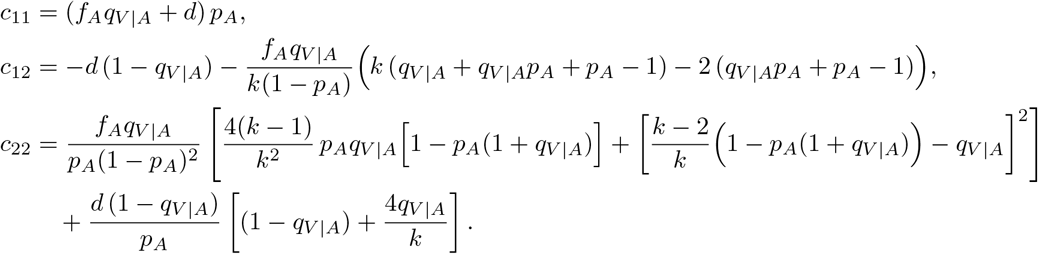

Taking the limit as *M* → ∞, Itô’s Lemma (Itô 1957) yields the SDE

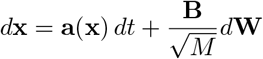

where

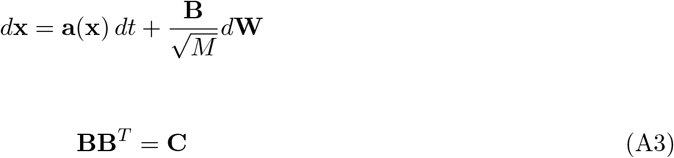

## B Analysis of the relaxed population distribution

### B.1 Carrying capacity for population structures

Based on the deterministic limit of the SDE equation for a *k*-regular graph (Equation 10), the pair-approximate carrying capacity of the resident population is

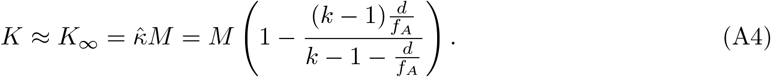

where 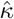 is the approximate equilibrium density defined in Equation 10. However, this derivation is based on the assumptions of pair approximation, which assumes independence of adjacent edges. Figure A1 shows that while the carrying capacity formula is accurate for both regular graphs and well-mixed graphs at low death rates, it loses that accuracy for higher death rates in the regular graph. At these higher death rates it overestimates the carrying capacity in the 3-regular graph. This overestimation occurs because increased clustering produces correlations between the states of neighbouring edges, violating the independence assumptions of the pair approximation. We focus on the 3-regular graph because these effects are expected to be most pronounced at low degree *k*.

**Fig. A1:**
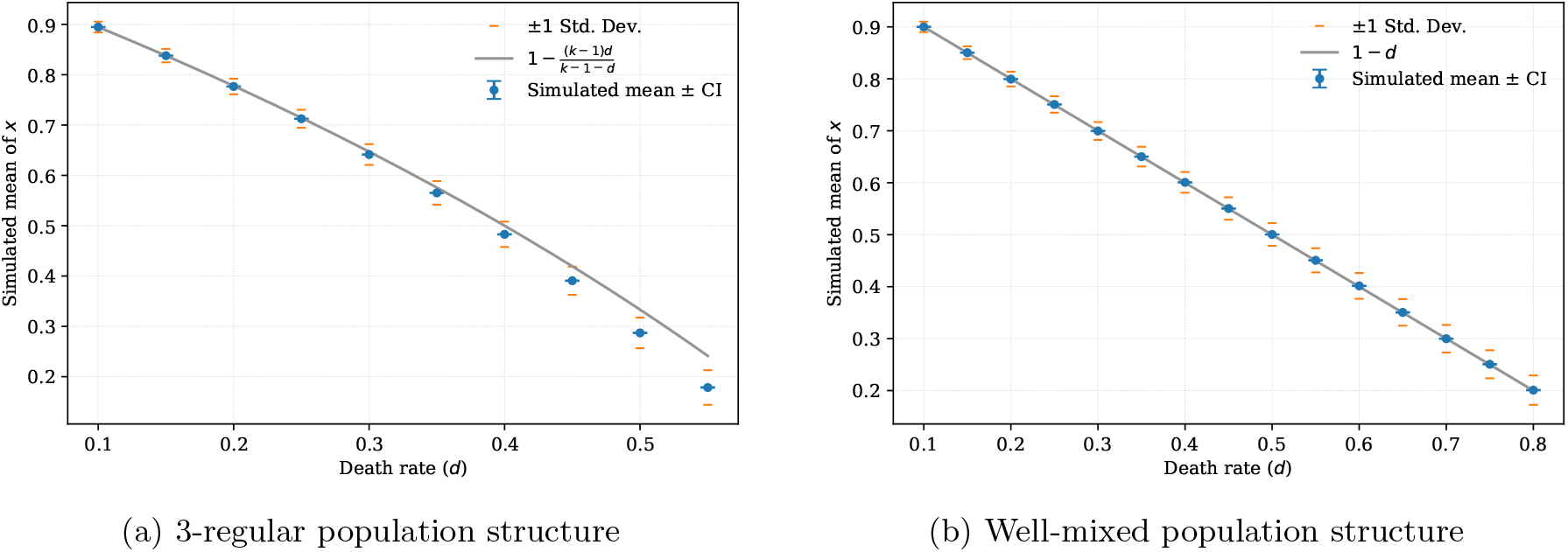
Mean resident population size (carrying capacity) as a function of death rate *d* for the decoupled single-type process, comparing simulations to deterministic predictions. Parameters are *f*_*A*_ = 1 and *M* = 1000. Points show simulation estimates and curves show the predicted carrying capacity *K*_∞_ (pair approximation for the regular graph and mean-field for the well-mixed case). Both structures show decreasing means and increasing standard deviations as the death rate increases. The well-mixed population’s mean-field approximation remains accurate throughout, but the pair approximation’s carrying capacity overestimates the mean population in the 3-regular graph, with the size of overestimation growing as the death rate increases.

As shown in Matsuda et al. (1992), pair approximations can be expanded to triple and higher order which produce more accurate estimates of the carrying capacity. Importantly, this reduces the maximal death rate at which a graph can sustain a population - when the carrying capacity is above 0. Based on pair approximation and a resident population with fitness 1, this death rate threshold value is 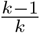, so we would expect this to be lower in reality.

### B.2 Effects of unsustainable death rates

Increasing the death rate past the point at which the population can be at quasi-equilibrium results in different takeover probabilities. An upper bound for the maximum death rate at which a regular graph can sustain a population can be found using the pair-approximate expression for *K* (Equation 10):

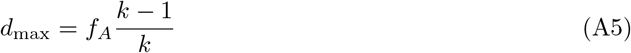

As the death rate increases towards *d*_*max*_, the neutral drift component from the death rate dominates. Consequently, the amplification effect in regular graphs transitions to a suppression effect for sufficiently weak selection, as shown for the 3-regular graph in Figure A2. For intermediate fitness values the amplification effect returns as at these fitness values the mutant population can be sustained, before transitioning again to suppression for sufficiently large *r*. Though this death rate is below *d*_max_ for *k* = 3, Equation A5 is based on pair approximation and we would expect the true threshold to be lower (Matsuda et al. 1992).

**Fig. A2:**
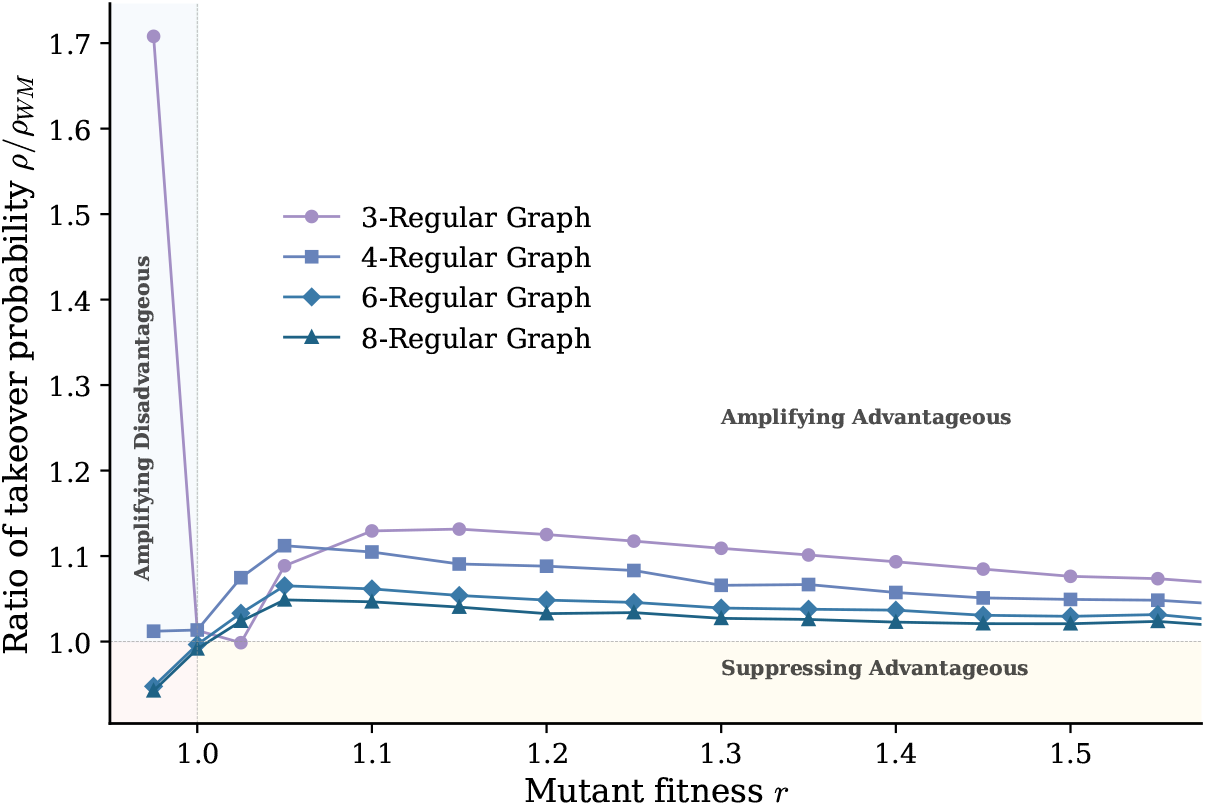
Ratio between takeover probabilities *ρ/ρ*_*W M*_ for the decoupled process on random regular graphs, comparing degrees *k* ∈ {3, 4, 6, 8} to the well-mixed baseline. Graph sizes *M* are chosen to approximately set *K* = 100 with *d* = 0.6: *M*_*W M*_ = 250, *M*_*k*=3_ = 1846, *M*_*k*=4_ = 431, *M*_*k*=6_ = 320, *M*_*k*=8_ = 295. Each point is computed from 10^6^ samples. The 3-regular graph shows an extreme ratio of *ρ/ρ*_*W M*_ ≈ 1.71 just below *r* = 1, while the higher-degree graphs remain close to 1. Just past *r* = 1, all curves jump to amplification, peaking near *r* ≈ 1.10–1.15 with peak height decreasing with degree, followed by a gradual decline as *r* increases.

### B.3 Equilibrium density ratio of regular graphs through pair approximation

The (pair-)approximation for the equilibrium density of a population with fitness *r* on a *k*-regular graph from Equation 10 is

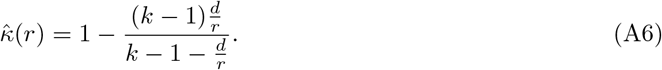

Under weak selection, *r* ≈ 1 + *w* where |*w*| ≪ 1. We explore how the equilibrium density changes on regular graphs using 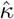:

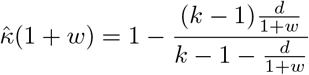

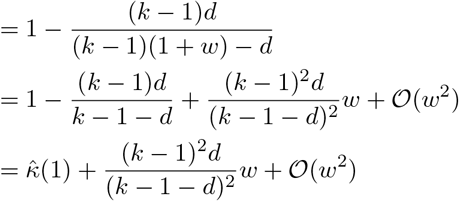

We can approximate the equilibrium density ratio using these quantities:

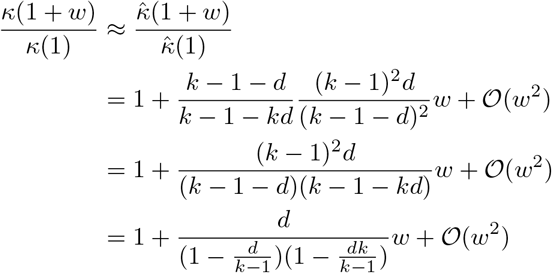

For weak selection, when |*w*| ≪ 1 the slope of the equilibrium density ratio is 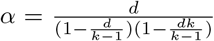.

For *k >* 3 and 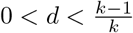, *α* increases with *d* and decreases with *k*.

### B.4 Effects of heterogeneous degree distribution

The degree distribution of a scale-free graph follows a power law, *P* (*k*) ~ *k*^−*γ*^, with exponent *γ* = 3 for the Barabási–Albert model. On this graph the relaxed resident population is no longer uniform across nodes: higher-degree nodes are more densely occupied than lower-degree ones, because a node with many neighbours is replaced more quickly after a death. This effect is stronger for larger death rates and lower average degree (Figure A3).

For low death rates when few vacant sites exist this results in suppression of selection, as the vacant space is disconnected (Figure 12). As the death rate increases, only the highest degree nodes remain occupied, allowing for vacant space to become more connected and for the mutant to reproduce and grow. This has less impact when ⟨*k*⟩ = 3 as many vacant nodes are leafs which are often disconnected from other vacant nodes.

**Fig. A3:**
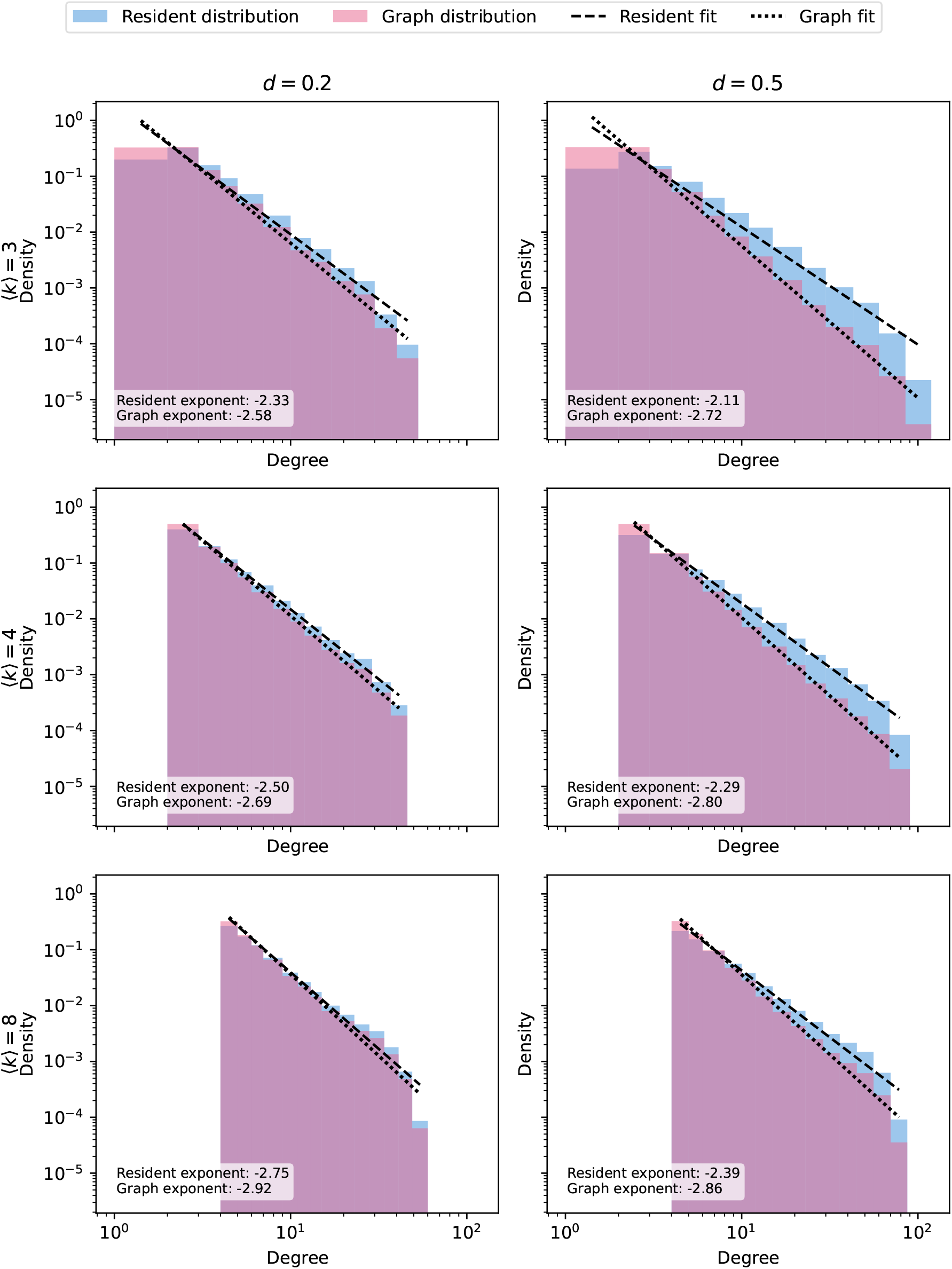
Histograms of node occupancy by degree in a relaxed resident population on different average degree Barabási-Albert graphs, normalised over their distributions. Lines of best fit overlay the histograms with exponents shown. The graph exponent tends to −3 as *M* → ∞(Barabási and Albert 1999). For all graphs, *f*_*A*_ = 1, *K* = 100. For *d* = 0.2, *M*_⟨*k*⟩=3_ = 187, *M*_⟨*k*⟩=4_ = 160, *M*_⟨*k*⟩=6_ = 148, *M*_⟨*k*⟩=8_ = 143. For *d* = 0.5, *M*_⟨*k*⟩=3_ = 811, *M*_⟨*k*⟩=4_ = 489, *M*_⟨*k*⟩=6_ = 355, *M*_⟨*k*⟩=8_ = 313. In all cases, the resident population is more concentrated in nodes of higher degree than the graph distribution.

## C Extinction time plots

Figure A4 shows the distribution of extinction times over 1000 simulations. As this is expected to be exponential (Ashcroft et al. 2015), the reciprocal of the mean extinction time is used as the maximum likelihood estimator of the rate of extinction.

**Fig. A4:**
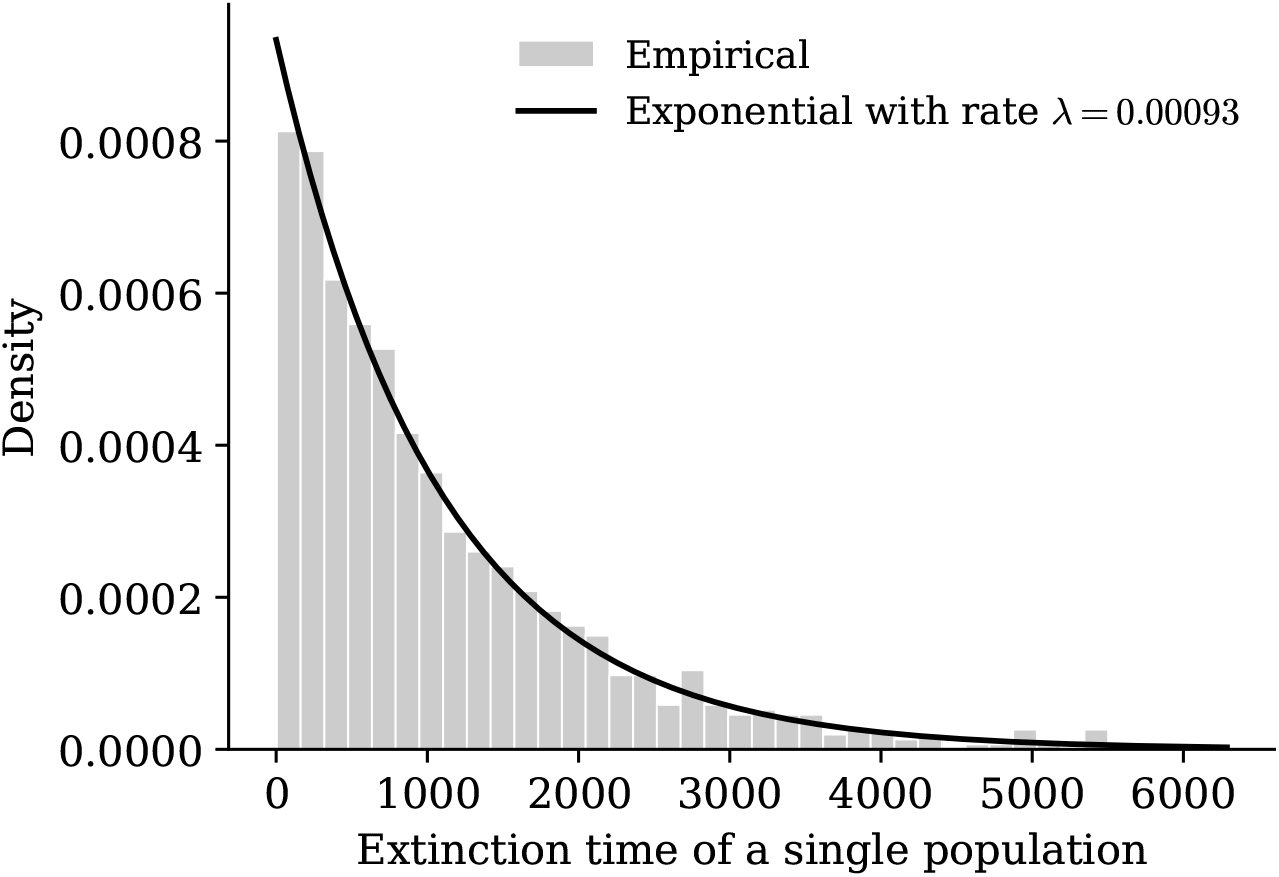
Distribution of extinction times with fitted exponential for the single type well-mixed population with *d* = 0.5, *f*_*A*_ = 1, *M* = 30 and 1000 simulations, starting at carrying capacity of *K* = 15. The extinction time distribution fits closely with the exponential distribution with rate 0.00093.

## D Lattice takeover probability for *d* = 0.2

In the main text, the takeover probability on the von Neumann and Moore lattices for a death rate of *d* = 0.5 is presented (Figure 11). Figure A5 shows the same comparison at *d* = 0.2. The qualitative behaviour is the same, with amplification just above *r* = 1 followed by suppression at large *r*, but the amplification of the lattices is closer to that of random regular graphs of the same degree, consistent with the smaller difference in expected density ratio slope at this death rate (Figure 10a).

**Fig. A5:**
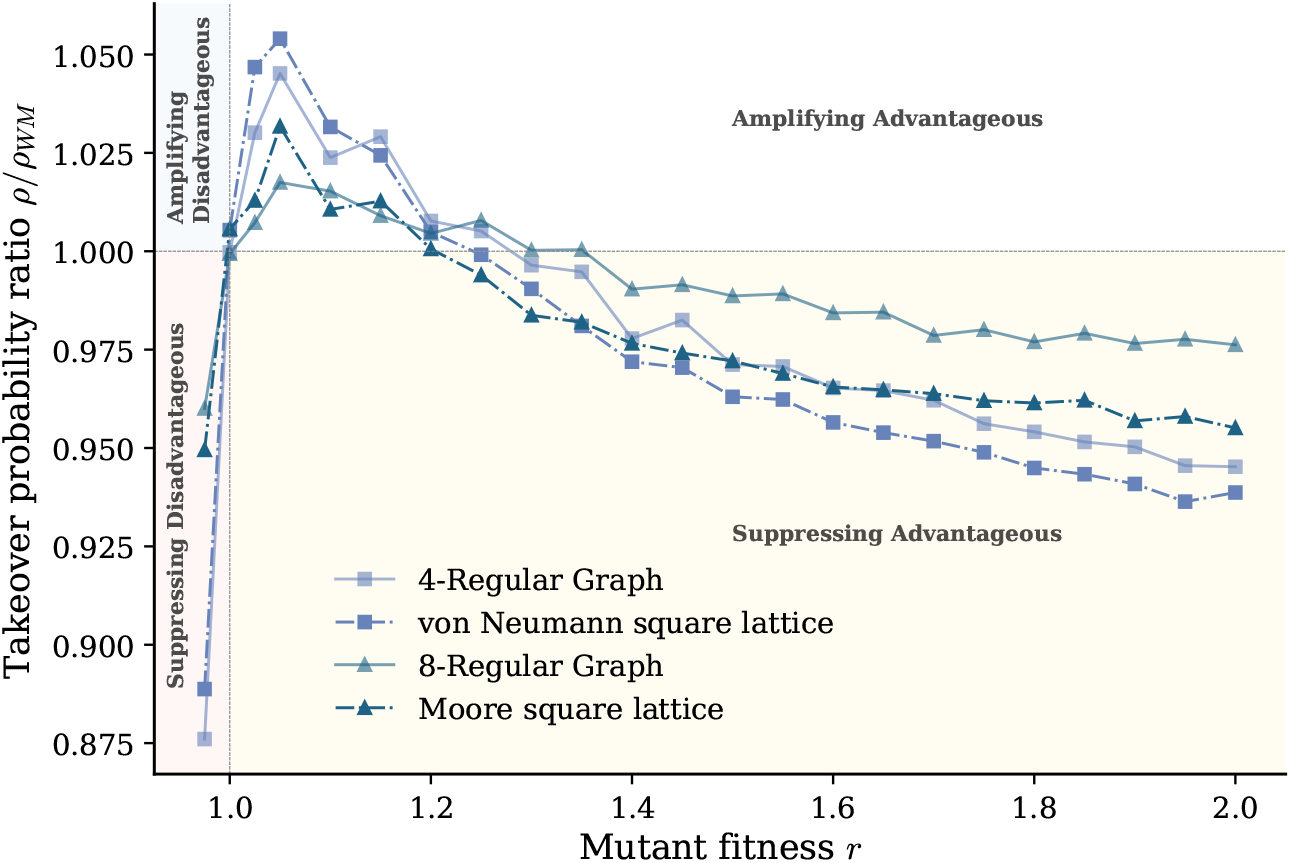
Ratio between takeover probabilities *ρ/ρ*_*W M*_ for the decoupled process on regular lattices, compared to the well-mixed baseline and random regular graphs of the same degree. Parameters are *d* = 0.2 with *M* = 121 (von Neumann lattice) and *M* = 121 (Moore lattice), and *M*_*k*=4_ = 128 and *M*_*k*=8_ = 126 for the corresponding random regular graphs. Each point is computed from 10^6^ samples. All four structures show a ratio below 1 for *r <* 1, a sharp jump to amplification just above *r* = 1, and a peak near *r* ≈ 1.05–1.10 before declining. This decline continues past *ρ/ρ*_*W M*_ = 1 into sustained suppression from *r* ≈ 1.3 onward. Each lattice initially amplifies more strongly than its corresponding random regular graph, but declines faster and falls below it once suppression sets in, after which the two decline in near-parallel fashion. Lower-degree structures amplify and suppress more than the higher-degree structures.

## E Connections between the Moran process and the decoupled process

### E.1 death-Birth: Limit as *d →* 0

The takeover probabilities for the decoupled process appear to tend to the fixation probability of the dB process as *d* → 0 (Figure 7). To make this thorough, we introduce the following theorem.

#### Theorem 1

*In the decoupled model, as the death rate tends to 0 the discrete Markov chain tends to the same Markov chain for the death-Birth coupled Moran model*.

*Proof* Consider a population at state *S* (population size *i* and some structure) in a space of size *M*. If the size is full such that *i* = *M* the next event must be death, so to prove the statement we must show that the probability of the next event being a birth for 0 *< i < M* individuals tends to 1. Let *p*_*B*_ (*S*) be the probability that a birth is successful and occurs on an vacant node. Then

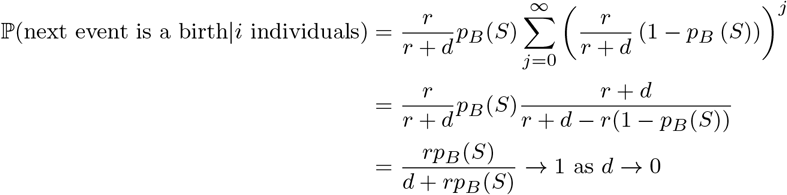

Therefore any population will increase to the maximum size and then alternate between *M* and *M −* 1. In a network population structure individuals will only be able to birth onto the singular vacant node if they are adjacent, which is equivalent to the death-Birth process. For populations with multiple types of varied fitness, *r* can be replaced with an average fitness and the same result is derived. □

### E.2 Birth-death: Well-mixed Upper Bound

For large *r* the takeover probabilities for all graphs decrease below the well-mixed takeover probability (Figure 8a, Figure 12a). Additionally, for these values of *r* the takeover probability of the well-mixed graph in the decoupled process is less than the fixation probability of the well-mixed graph for the coupled Moran process (Figure 6). We formalise these concepts in the following theorems.

#### Theorem 2

*In the decoupled model, for sufficiently large fitness ratio r and sufficiently large carrying capacity K, the takeover probability of the decoupled process for any graph with bounded degree is bounded above by the well-mixed graph*.

*Proof* We consider the takeover probability as *r → ∞*. In this limit the probability that the next event being a mutant reproducing tends to 1 if it is adjacent to a vacant node.

For all graphs, if the mutant reproduces it cannot go extinct: if either of the two (or more) mutants dies then a vacant node is produced that is adjacent to a mutant. The adjacent mutant will then reproduce onto it with probability *→* 1 as *r → ∞*. Because of this we can view the mutant as taking over as soon as it reproduces once.

For a well-mixed population/complete graph of size *M* with carrying capacity *K*, the mutant will be adjacent to a vacant node when it is introduced unless the resident population size is *M −* 1. If the resident population size is *M −* 1 then the mutant goes extinct with probability 1*/M*, otherwise a resident dies and an adjacent vacant node is created. Let *ρ*_W-M_ be the takeover probability for a mutant in a well-mixed population in the limit *r → ∞*. Then

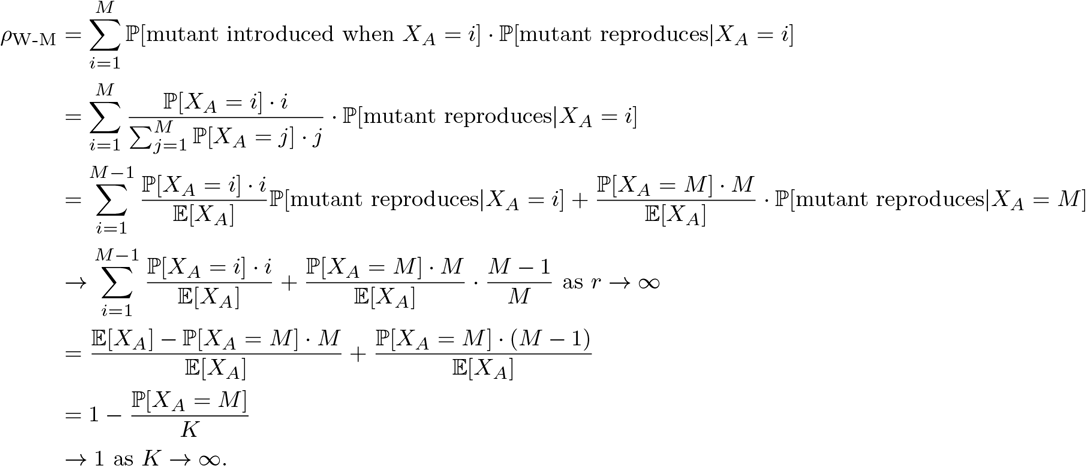

Consider now a graph *G* of size *M*_*G*_ with max degree *k*_*max*_ *< M*_*G*_. Let *k*_*j*_ be the degree of node *j*. Then

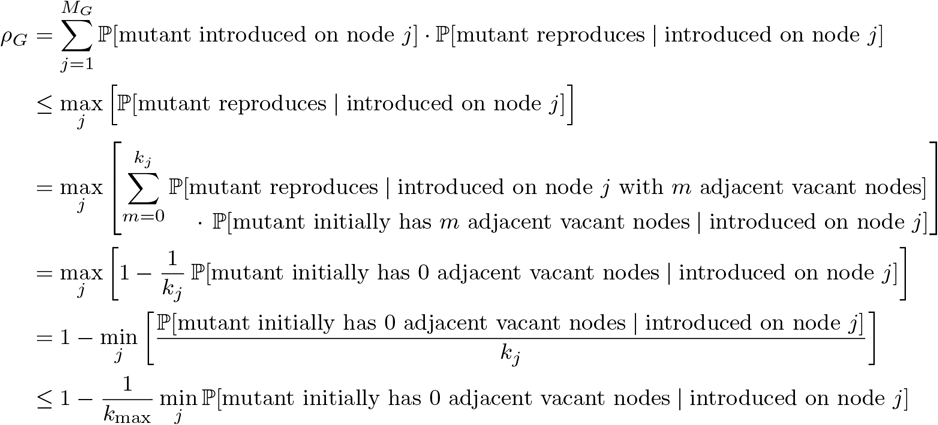

We show that this value is strictly less than 1. Consider an occupied node *j* and its adjacent neighbours, forming a star. Independent of the rest of the graph, an adjacent vacant neighbour has a rate of *f*_*A*_*/k*_*j*_ of becoming occupied by node *j* reproducing. This means that the probability of a neighbour being occupied is strictly greater than 0. As the occupancy rate is only increased by other nodes in the graph, this probability can only increase with *M*_*G*_. Therefore the probability that a mutant has 0 adjacent vacant nodes is bounded below by some *c*(*k*_*j*_, *f*_*A*_, *d*) *>* 0.

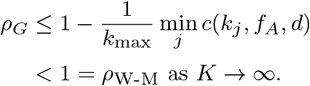

□

This result is consistent with our simulations: for random regular graphs, the ratio *ρ/ρ*_WM_ of the graph takeover probability to the well-mixed takeover probability falls below 1 and converges to a constant as *r* increases (Figure A6).

**Fig. A6:**
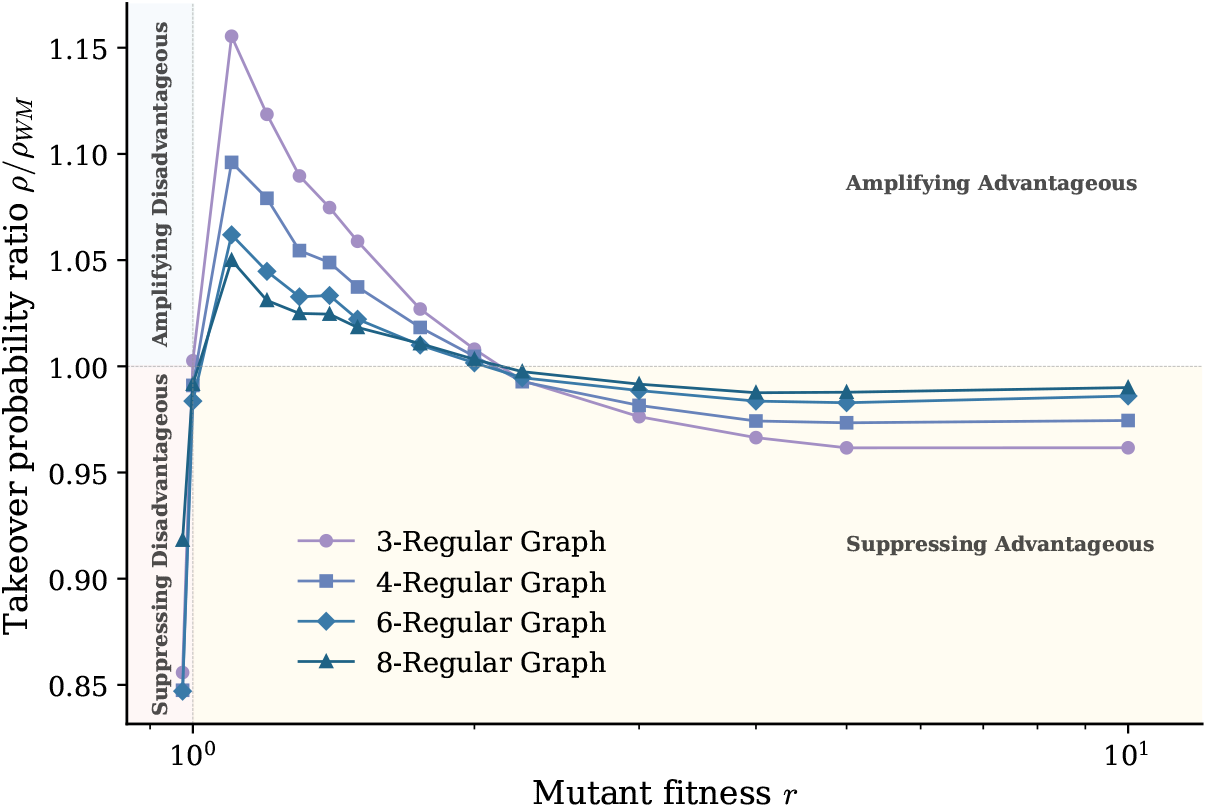
Ratio between takeover probabilities *ρ/ρ*_*W M*_ for the decoupled process on random regular graphs, comparing degrees *k* ∈ {3, 4, 6, 8} to the well-mixed baseline. Graph sizes *M* are chosen to approximately set *K* = 100 with *d* = 0.5: *M*_*W M*_ = 200, *M*_*k*=3_ = 341, *M*_*k*=4_ = 259, *M*_*k*=6_ = 230, *M*_*k*=8_ = 219. Each point is computed from 10^6^ samples. The ratio *ρ/ρ*_WM_ is below 1 for *r <* 1, then rises sharply, crossing 1 at *r* = 1 and reaching amplification just above it. For every graph degree it peaks at *r* ≈ 1.05–1.15, with lower peaks at higher degree, and then declines gradually as *r* increases. The curves for different degrees are approximately equal at *r* = 2, and beyond that they continue to decrease, with the largest decrease for the lowest degree. By *r* = 10 the ratio has settled to a constant value for each degree.

#### Theorem 3

*For sufficiently large fitness r the takeover probability in the decoupled process on a well-mixed population is bounded above by the fixation probability of the coupled Birth-death process in the well-mixed population. As the carrying capacity K additionally tends to ∞, the takeover probability tends to 1 – the fixation probability of the Bd process*.

*Proof* The fixation probability of the Birth-death process in the well-mixed population is

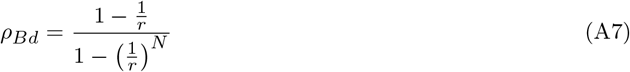

for system size *N*. This has limit 1 as *r → ∞*. The takeover probability of the decoupled process in the well-mixed population in the limit *r → ∞* is

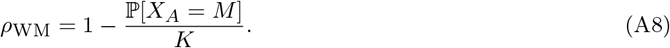

This value is less than 1, so *ρ*_*Bd*_ *> ρ*_WM_ as *r → ∞*. As *K → ∞, ρ*_WM_ *→* 1 and the result follows. □

